# Sugar-mediated control of *ATG101* modulates carbon deficiency-induced autophagy in Arabidopsis

**DOI:** 10.1101/2025.01.12.632587

**Authors:** Svetlana Boycheva Woltering, Mirai Tanigawa, Tobias Bläske, Erika Isono

## Abstract

Autophagy is a major eukaryotic degradation and recycling pathway, which eliminates damaged cellular components and provides energy and building blocks especially under adverse environmental conditions. While the core autophagy machinery has been extensively studied, autophagy initiation and regulation remain less understood. In this study, we show that genes encoding the core autophagy initiation complex, *ATG1*, *ATG11*, *ATG13*, and *ATG101*, are transcriptionally regulated by sugars. To analyze the expression of these genes, we took advantage of *Arabidopsis thaliana* accessions, which display strong variations in their responses to darkness-induced fixed carbon deprivation. One of the components of the initiation complex, *ATG101*, has multiple sugar-related elements, and was repressed by sugar. Moreover, *ATG101* was induced stronger upon starvation in carbon starvation-resistant accessions when compared to carbon starvation-sensitive accessions. We identified three single nucleotide polymorphisms in the genomic region of *ATG101* which are associated with the carbon-starvation phenotype. Further analyses through complementation of the *atg101* mutant with different variants of *ATG101* demonstrated that the single nucleotide polymorphism alleles contribute differently to *ATG101* expression regulation, response to sugar, and phenotype recovery. Taken together, our data suggest that modulation of *ATG101* by sugar is part of a multi-layered mechanism regulating the sensitivity to carbon starvation in Arabidopsis.

## Introduction

Plants employ several recycling pathways to optimize energy consumption and recycle nutrients, as a way to withstand unfavorable environmental conditions and ensure their own survival. One of those catabolic routes is (macro)autophagy, a highly conserved mechanism which consists of enclosing cytoplasmic material followed by its delivery to the vacuole for degradation (Li and Vierstra, 2012; Liu and Bassham, 2012). The process involves the formation of a cup-shaped structure, the phagophore. As autophagy progresses, the phagophore expands and eventually closes, resulting in a double-membrane vesicle, the autophagosome. The latter is subsequently delivered and fused to the vacuole, followed by degradation of the autophagosomal content.

Autophagy initiation and progression involve a system of AUTOPHAGY (ATG) proteins which are highly conserved in eukaryotes although certain variation exists among the kingdoms. *ATG* genes were first discovered and characterized in yeast (Tsukada and Ohsumi, 1993) and have now been studied in many organisms from all lineages, including plants (Tsukada and Ohsumi, 1993; Liu and Bassham, 2012). The core autophagy machinery can be divided in five groups of proteins the first one of which is the initiation complex of ATG1 protein kinase. The second protein complex comprising ATG9/ATG2/ATG18 is involved in delivery of membrane material for the growing phagophore while the phosphatidyl inositol-3-kinase (PI3K) complex, consisting of ATG14, VACUOLAR PROTEIN SORTING (VPS)15 and VPS34, is responsible for vesicle nucleation. The fourth group of proteins includes the two ubiquitin-like conjugation systems ATG8-phosphatidyl ethanolamine (PE) and ATG5-ATG12, which regulate autophagosome expansion (Thompson et al., 2005). The fifth and last group contains the soluble N-ethylmaleimide sensitive factor (NSF) attachment protein receptor (SNARE) and participates in the fusion of the autophagosome with the vacuole (Liu and Bassham, 2012; Marshall and Vierstra, 2018).

Currently, good understanding exists of the molecular function of the core autophagic machinery, while regulation of the autophagic activity remains insufficiently explored. A large part of the natural accessions of Arabidopsis has been investigated in terms of single nucleotide polymorphisms (SNPs) (Kim et al., 2007; Zeller et al., 2008; Zhang and Borevitz, 2009) and/or re-sequenced (Nordborg et al., 2005; Alonso-Blanco et al., 2016) making them a valuable tool to explore the relationship between genetic variation and phenotype, and its contribution to adaptation and phenotypic plasticity. As these Arabidopsis variants, or accessions, originate from a variety of habitats, they are adapted to diverse environmental conditions. Therefore, differences in autophagy induction, efficiency and regulation due to diversified responses could be expected for these accessions.

Autophagy is a constitutive process that maintains the homeostasis of the cell (Masclaux-Daubresse et al., 2017), and occurs constantly at low levels (Marshall and Vierstra, 2018), which can be activated by various types of abiotic and biotic stress. Nutrient deficiencies such as nitrogen or carbon starvation can trigger autophagy. It has been established that inhibition of the TOR kinase by nutrient deficiency, such as fixed carbon starvation due to prolonged darkness (hereafter referred to as carbon starvation), can lead to autophagy activation. The TOR kinase is activated by sugar, repressed in its absence and is itself regulating most responses governed by sugar (Wullschleger et al., 2006; Dobrenel et al., 2013; Xiong and Sheen, 2015). It is however unclear, if sugar is regulating the expression of the downstream autophagy genes. Typically, under optimal growth conditions, the target of rapamycin (TOR) kinase phosphorylates ATG13 and thus prevents the initiation of autophagy by inhibiting the complex formation between ATG1 and ATG13 (Liu and Bassham, 2010; Puente et al., 2016). Stress conditions such as nutrient deficits inhibit TOR and lead to the ATG13 dephosphorylation and assembly of the autophagy initiation complex together with ATG1, ATG11 (RB1CC1/FIP200), and other accessory proteins which vary between the different lineages. For instance, most eukaryotes, including plants and mammals, possess ATG101 but lack ATG17, ATG29 and ATG31, that are found in *Saccharomyces cerevisiae* (Alers et al., 2014; Noda and Mizushima, 2016). The role of ATG101 is to stabilize ATG13 and maintain its complex with the kinase ATG1 as described by Mercer and co-workers (Mercer et al., 2009) and thus, it is essential for autophagy (Mercer et al., 2009; Qi et al., 2015; Suzuki et al., 2015; Kim et al., 2018).

The structure of the ATG13-ATG101 complex has been shown to be a Hop1, Rev7, Mad2 (HORMA) domain heterodimer (Qi et al., 2015; Suzuki et al., 2015), suggesting it could coordinate the regulation of autophagy induction by orchestrating the binding of substrates and regulatory proteins. Although the *A. thaliana* ATG101 has only 26 % homology with its mammalian counterpart, it interacts with ATG13 and ATG11 (Li et al., 2014), like the mammalian homolog. *A. thaliana atg101* knockout mutant plants show increased sensitivity to prolonged darkness (Lee et al., 2023).

In this study, we have identified sugar related cis-regulatory elements in the promoters and introns of genes encoding autophagy initiation complex components *(ATG1a, ATG1b, ATG1c, ATG11, ATG13a, ATG13b,* and *ATG101*) and determined their response to soluble sugars. More specifically, we have compared the response to sucrose with the effect of glucose, galactose and arabinose, in addition to the sugar alcohol mannitol. All investigated genes were found to be suppressed by sugar treatment but not by mannitol.

Additionally, we performed an assessment of a set of 181 *A. thaliana* accessions, originating from different parts of the world. They were evaluated in terms of their sensitivity to carbon starvation (darkness-mediated) by measuring the loss of chlorophyll (darkness/light ratio). As a result, we were able to establish carbon starvation-resistant (CS^R^) and carbon starvation-sensitive (CS^S^) accessions. Subsequently, we focused on a set of CS^R^ and CS^S^ accessions and found that among the initiation complex genes, only *ATG101* is differentially regulated in the investigated CS^R^ and CS^S^ accessions. Nitrogen deprivation did not result in the same chlorophyll ratio pattern and neither did it cause any significant induction of *ATG101* gene expression.

We identified three separate single nucleotide polymorphisms (SNPs) in the *ATG101* gene that correlate with the observed carbon starvation phenotype. The two main alleles of those SNPs associated strongly with the resistant and sensitive phenotype of the accessions. Complementing the *atg101* knockout mutant with the different alleles demonstrated that only the resistant variant could fully rescue the mutant phenotype. In summary, our findings suggest that sugar regulates the expression of *ATG101* gene. The gene exhibits variations across natural accessions linked to the presence of specific SNPs in the genomic region. Thus, *ATG101*, through its interplay with sugar, could directly contribute to the regulation of darkness-induced autophagy.

## Results

### Expression of ATG1-ATG13 complex encoding genes is modulated by sugar

Sugar has been established as a signaling molecule in autophagy, with different sugar species regulating the process under nutrient deficiency (Rolland et al., 2002; Janse van Rensburg et al., 2019). Responses to sugar at the transcriptional level can be mediated by cis-regulatory elements. While it is clear that abundance of various sugars generally suppresses autophagy (Janse van Rensburg et al., 2019), little is known about the transcriptional regulation of autophagic genes in response to sugar. We reasoned that sugar could affect the abundance and/or activity of upstream components regulating nutrient deficiency-triggered autophagy. We focused on the ATG1-ATG13 initiation complex, whose assembly is essential for downstream events during autophagy. We first wanted to establish whether genes encoding subunits of the ATG1-ATG13, namely *ATG1a, ATG1b, ATG1c, ATG11, ATG13a, ATG13b*, and *ATG101*, have sugar-related cis-elements in the promoters (- 1500 bp upstream of the transcription start site) or introns. We obtained all the sequences from phytozome.org and looked for the following set of elements: CGACG, TATCCA and ACGTG, TTATCC, TATCCAY, CATCC, ATATCT and GATAA, GGATA, and GATTA previously implicated in sugar perception or overrepresented in promoters of sugar suppressed genes such as the *Sugar Transporter Protein (STP) 1* and the *Dark-Induced 6 (DIN6)* gene as summarized by Cordoba and colleagues (Cordoba et al., 2015).

The presence of these elements leads to repression of gene expression upon addition of sugar, except for CGACG which confers gene activation during sugar starvation. *ATG101* contains all ten elements while the rest of the investigated genes have a subset of these elements in different combinations (Figure 1A). Next, we tested in 1-week-old Col-0 seedlings if *ATG1a, ATG1b, ATG1c, ATG11, ATG13a, ATG13b*, and *ATG101* actually respond to a sugar treatment. We have chosen sucrose and glucose as two of the most abundant soluble sugar species in Arabidopsis (Ward et al., 1997; Ruan, 2014). In addition, we have treated seedlings with galactose and arabinose, two sugars found in the extracellular matrix of Arabidopsis (Zablackis et al., 1995; Gangl and Tenhaken, 2016) as well as mannitol to rule out osmotic stress response. All sugars suppressed the expression of the ATG1-ATG13 kinase complex genes for the used concentration and durations (Figure 1B, C, D and E). Mannitol, on the other hand, had no strong effect (1 hour treatment) or slightly induced the genes (2 h treatment; Figure 1F). We also looked at the nine *ATG8* homologs. ATG8 is a structural component of the phagophore and the autophagosome, acting downstream of the ATG1-ATG13 complex. Sugar-related cis-elements were abundant in all *ATG8* loci, except for *ATG8c,* in which only two types of elements could be found (GATTA and GATAA). We tested the response of the *ATG8c, ATG8e, and ATG8f* to sucrose and found that *ATG8e* and *ATG8f* which contain nine out of the ten elements were downregulated by the addition of sucrose, whereas *ATG8c* is not affected (Supplementary Figure S1), suggesting that not all autophagy-related genes are transcriptionally regulated by sugar.

**Figure 1.**
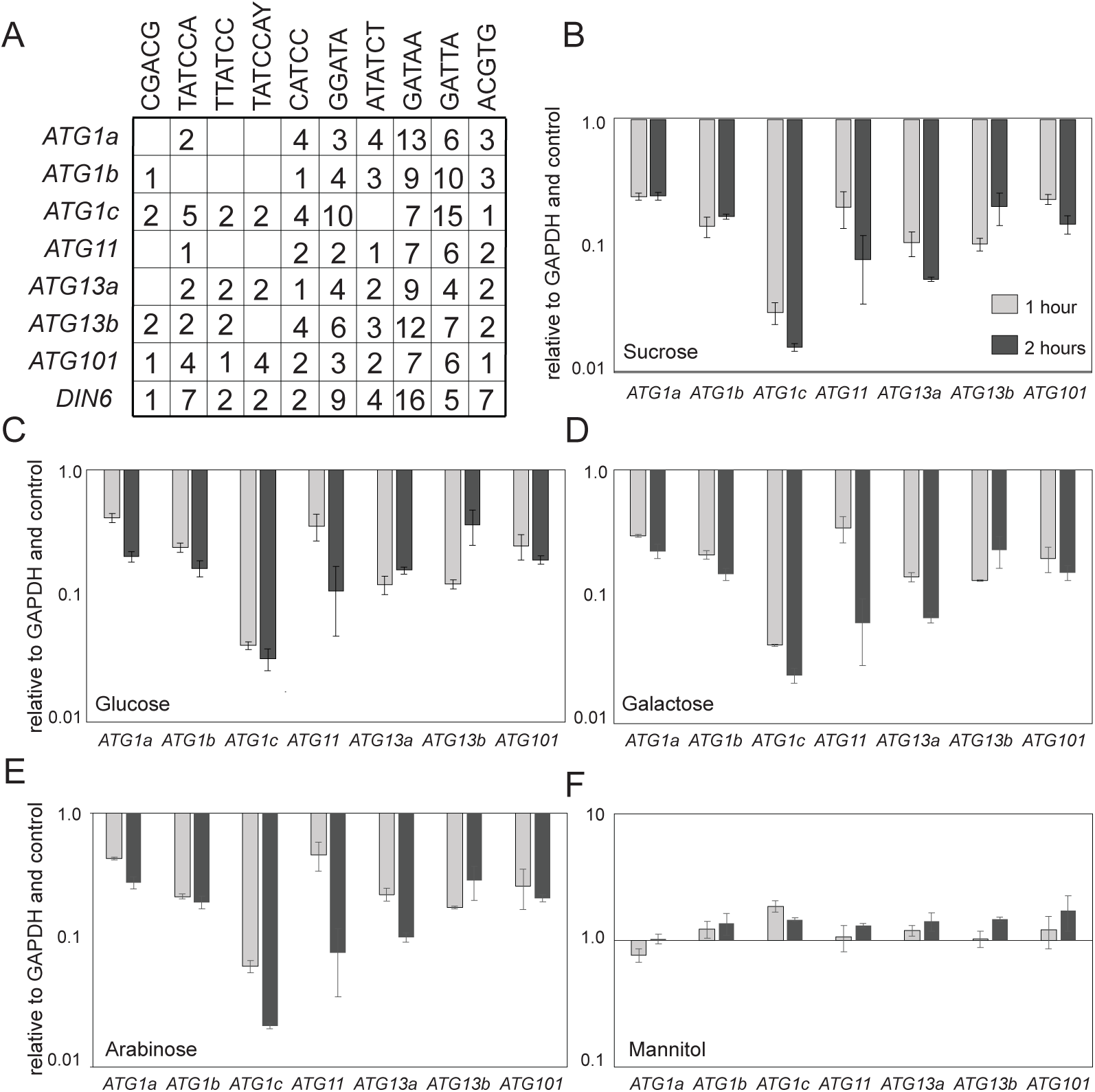
The expression of *ATG* genes encoding ATG1-ATG13 complex components is modulated by various sugars. A. Cis-regulatory elements related to sugar are found in the promoters (-1500 bp) and introns of *ATG* genes. B - F. Effect of 2% concentration of different sugars and mannitol on the expression of *ATG1* and *ATG13* homologues, *ATG11* and *ATG101.* Data represent mean values of two independent experiments with one biological and three technical replicates each; error bars represent standard error.B. Sucrose treatment. C. Glucose treatment. D. Galactose treatment. E. Arabinose treatment. F. Mannitol treatment.

### *A. thaliana* accessions respond differently to carbon starvation

In order to understand the impact of the identified sugar suppression of autophagy-related genes on plant responses to carbon starvation, we decided to utilize a collection of natural accessions of *A. thaliana.* Natural accessions of *A. thaliana* grow in diverse environmental conditions, and are therefore adapted to local sets of factors. The accessions would consequently respond differently to changes in those factors, including but not limited to, light quantity and quality, temperature, and soil conditions. Thus, it can be expected that the accessions react differently to nutrient deficiency-induced autophagy (Lasky et al., 2014; Kang et al., 2023).

In order to determine whether Arabidopsis accessions have different response to carbon starvation, we first evaluated the chlorophyll retention after prolonged dark treatment. For the experiment, seedlings from 181 different accessions were subjected to seven days of darkness (subset; Figure 2A). Chlorophyll content was measured and the ratio between dark-treated and control seedlings was calculated (50 accessions shown Figure 2B, remaining results Supplementary Figure S2). Chlorophyll ratio varied considerably among the accessions. Whereas some accessions lost most of their chlorophyll, others retained more than 90 % of it (Figure 2B upper panel and Supplementary Figure S2), suggesting that the sensitivity of Arabidopsis seedlings towards carbon starvation differs among the ecotypes.

**Figure 2.**
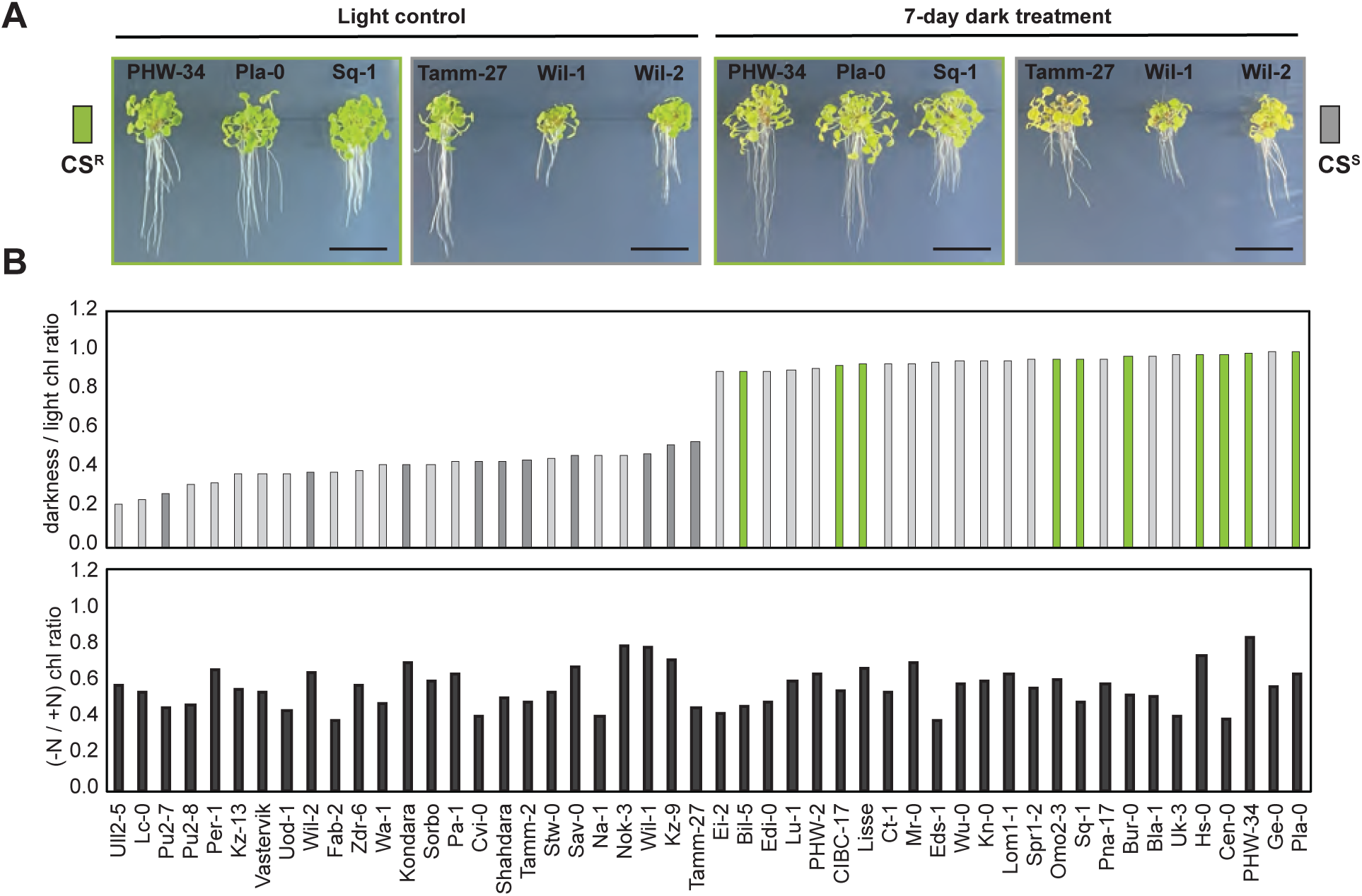
Darkness-induced fixed carbon starvation and nitrogen deprivation affect differently a range of *A. thaliana* accessions. A. Phenotypic changes in 1-week-old seedlings of six *A. thaliana* accessions after a 7-day dark treatment. B. Chlorophyll ratios: upper panel dark treated/light chlorophyll ratio of *A. thaliana* seedlings; lower panel nitrogen deficient/nitrogen containing chlorophyll ratio of *A. thaliana* seedlings.

To investigate whether this is a general starvation-effect, we next analyzed the chlorophyll content under nitrogen deficiency. For this purpose, we took 25 accessions from the ones with highest darkness to light chlorophyll ratio and 25 accessions among those with lowest values and subjected them to nitrogen starvation by transferring 1-week-old seedlings to nitrogen free medium for 2 weeks. Chlorophyll measurements of control and nitrogen starved plants showed a wide distribution of responses (Figure 2B, lower panel). The distribution, however, did not correlate with the pattern observed after carbon starvation with a Pearson correlation coefficient of 0.06 (p = 0.676), suggesting that different genetic basis determine the response of the seedlings to carbon-and nitrogen starvation.

### *ATG101* is differentially expressed in CS^R^ and CS^S^ accessions

We classified the accessions according to their carbon-starvation response. More specifically, we refer to accessions which retained more than 85 % of their chlorophyll as carbon starvation-resistant (CS^R^) and to those that retained less than 45 % of the control level as carbon starvation-sensitive (CS^S^). For further experiments we used the following eight ecotypes as representatives of each of the groups: CS^R^: Bur-0, Hs-0, Omo2-4, and Pla-0, and CS^S^: Cvi-0, Kondara, Kz-9, and Pu2-7 (Figure 3A).

**Figure 3.**
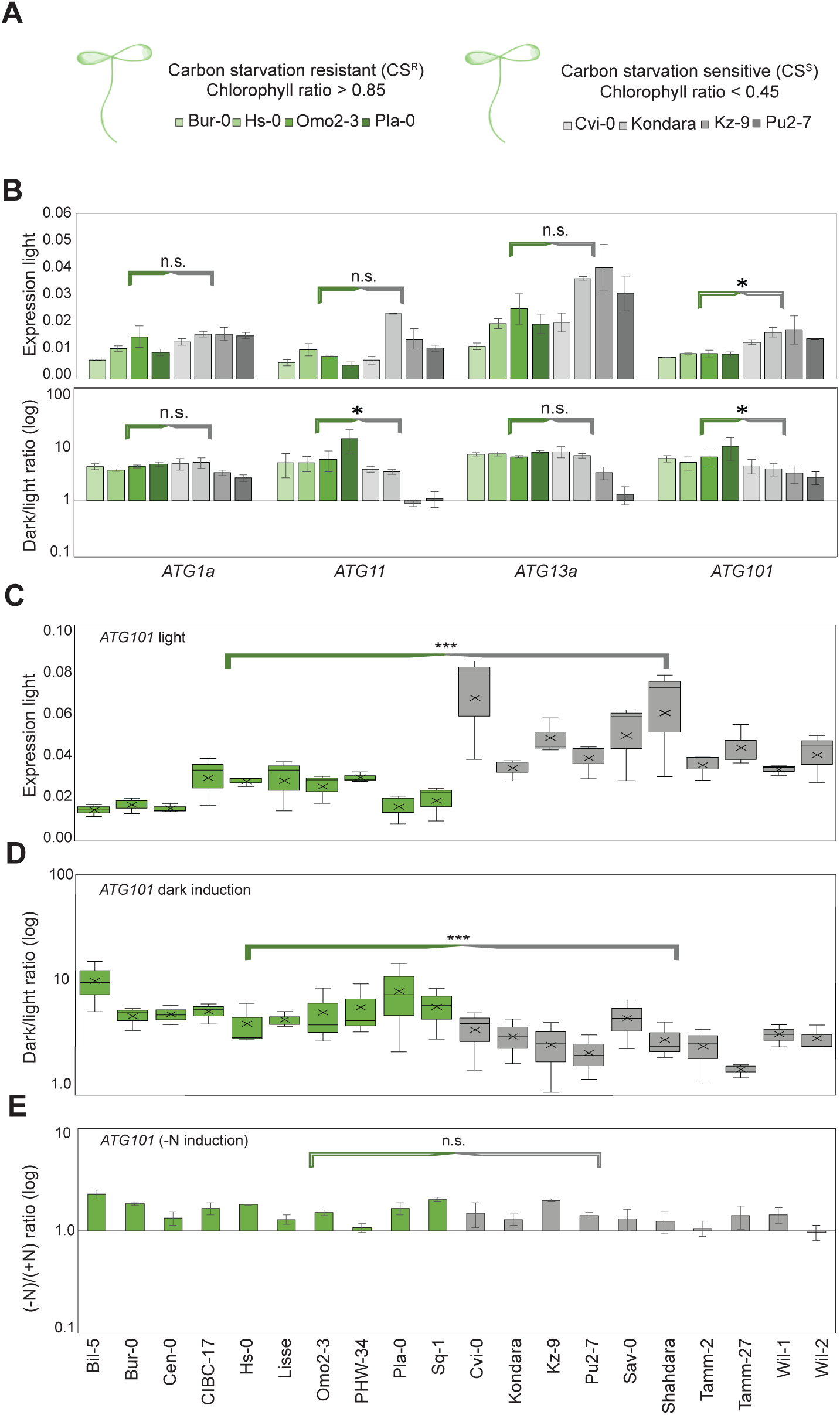
*ATG101* gene, encoding an autophagy initiation complex component is differentially regulated in carbon starvation-resistant and carbon starvation-sensitive accessions. A. Selected CS^R^ and CS^S^ accessions based on their dark-treated/light chlorophyll ratio. B. Upper panel: expression of *ATG1a*, *ATG11 ATG13a* and *ATG101* in CS^R^ and CS^S^ accessions under control conditions (light). Lower panel: expression of *ATG1a*, *ATG11, ATG13a* and *ATG101* in CS^R^ and CS^S^ accessions after 24h of dark treatment. All values are average of two independent experiments with one biological and three technical replicates each. Statistically significant differences were determined by two-tail t-test to compare the CS^R^ and CS^S^ groups and assuming unequal variance; * indicates statistically significant differences between the groups for p<0.05 while n.s. indicates the absence of such. C. Expression of *ATG101* in control seedlings of CS^R^ and CS^S^ accessions, in 20 *A. thaliana* accessions. D. Induction of *ATG101* gene by 24 h-darkness in 20 *A. thaliana* accessions. E. Changes in *ATG101* expression due to nitrogen deficit. Data in C and D are mean values from three independent experiments conducted with one biological and three technical replicates each. Error bars represent positive and negative variation, cross in each bar indicates the mean value. Data in E are mean from two independent experiments conducted with one biological and three technical replicates each. Error bars represent standard error of the means between the biological replicates involved. Statistics were performed using a two-way t-test assuming unequal variance. Levels of significance are indicated by asterisk (*** p < 0.0005) or n.s. in case no significance was found.

We next tested the possibility, that these accessions have differences in autophagy induction. We investigated the expression of *ATG1a, ATG11, ATG13a,* and *ATG101* genes. After a dark treatment, all genes were induced, however, significant differences between the two groups (CS^R^ and CS^S^) were observed only for *ATG101* with the CS^R^ lines displaying a higher level of gene activation (Figure 3B lower panel). Differences in the expression of those genes between the CS^R^ and CS^S^ group were also observed under control conditions where the resistant accessions exhibited lower expression (Figure 3B upper panel). We focused our further analyses on *ATG101* as it showed statistically significant differences between CS^R^ and CS^S^ accessions in both control and dark treatment conditions.

To corroborate this result, we repeated the analyses with 10 CS^R^ and 10 CS^S^ lines; CS^R^ – Bill-5, Bur-0, Cen-0, CIBC17, Hs-0, Lisse, Omo2-4, PHW-34, Pla-0 and Sq-1; CS^S^ – Cvi-0, Kondara, Kz-9, Pu2-7, Sav-0, Sha, Tamm-2, Tamm-27, Wil-1 and Wil-2). The obtained results confirmed that also in the extended analyses, CS^S^ accessions have altogether higher *ATG101* expression when grown in light and the gene is induced more strongly by carbon starvation in CS^R^ ecotypes (Figure 3C and 3D).

We next analyzed the expression level of *ATG101* after nitrogen starvation. In contrast to carbon starvation, the expression of *ATG101* was not substantially induced in any of the analyzed accessions after 48 h of (-N) treatment (Figure 3E) or 72 h of (-N) treatment (data not shown), suggesting that the ready induction of *ATG101* is a feature of carbon-starvation.

### TOR kinase activity is suppressed more strongly in CS^R^ than in CS^S^ accessions

The assembly and functioning of the ATG101-containing autophagy initiation complex depends directly on the inactivation of the TOR kinase (Liu and Bassham, 2010). Thus, probing the activity of TOR in CS^R^ and CS^S^ accessions would allow us to examine the events upstream of the ATG1-ATG13 complex. The TOR kinase activity was measured as the ratio of phosphorylated (P) versus unphosphorylated (NP) RPS6 (Figure 4A and B).

**Figure 4.**
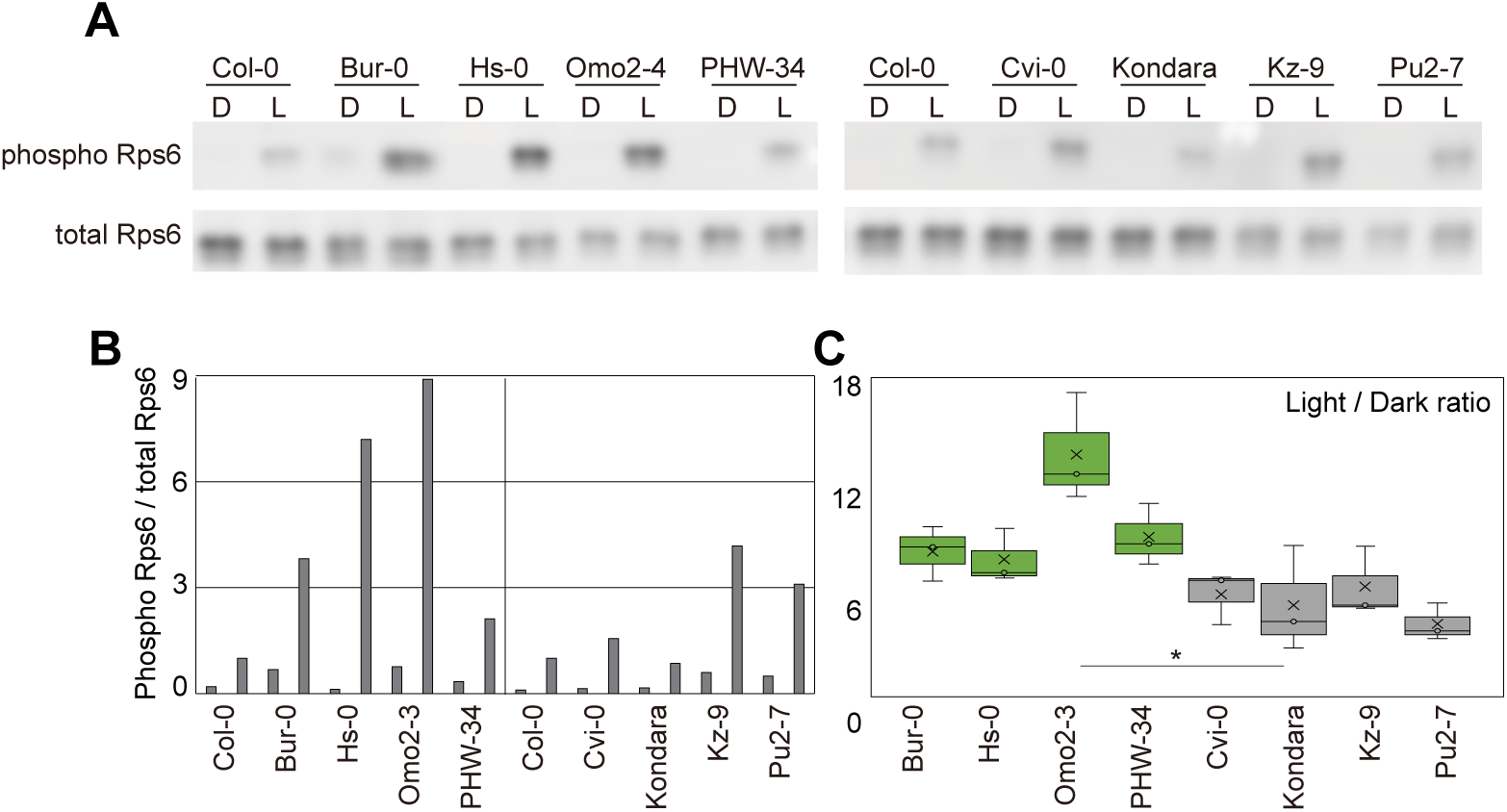
TOR kinase activity shows variation between darkness resistant and darkness sensitive accessions. A. Western blot analysis using an anti-Rps6 and an anti-phosphorylated Rps6. Rps6 is a secondary TOR kinase target and after quantifying the bands on the Western blots, the ratio of phosphorylated vs. total Rps6 protein levels in light grown and dark treated seedlings were determined. Eight *A. thaliana* accessions were used. The experiment was conducted three times and one representative result is shown. B. Quantification analysis of the WB results form A. C. Box plot of the ratio between TOR activity in light-grown and dark-treated seedlings. For C, three independent experiments are depicted by the box plots, prior to which values were normalized against the respective Col-0 values. Statistically significant differences between the two groups were detected for p = 0.044 (*; based on a two-way t-test analysis assuming unequal variance).

RPS6 is a second-degree target of TOR, and is phosphorylated by the ribosomal S6 kinase (Shi et al., 2018). The ratio between TOR kinase activity of control and dark treated seedlings displayed significantly higher levels in CS^R^ than in CS^S^ accessions (Figure 4C; p = 0.044). Thus, in the eight accessions examined, there are significant differences in the TOR kinase activity suppression by darkness between the CS^R^-and CS^S^ accessions. No such patterns in the TOR activity when control and dark treated seedlings were analyzed separately (Sup. Figure S3A to S3B). It is therefore likely, that the differences between TOR kinase activity in light and dark-treated plants might be important for autophagy activation and by extension resistance to darkness.

### Carbon starvation resistance/sensitivity patterns correlate with the main allelic combinations of SNPs in *ATG101* locus

To understand the genetic basis for different sensitivity patterns of the accessions, we investigated SNPs situated in the *ATG101* locus for all 181 accessions (https://gwas.gmi.oeaw.ac.at; Figure 5A). Out of the 17 SNPs with minor allele frequency of at least 15 (Figure 5A; lower panel), only three had a p-value of 10^-5^ or lower when correlated with the chlorophyll ratio. The first of those SNPs was in the second intron [T/C (Chr5: 26726206], the second lead to a synonymous substitution [T to A (Chr5: 26726627)], while the third was also intronic, resulting in a C/A (Chr5: 26726846) change. In the 181 accessions that we analyzed, C was more frequent than T in the first SNP, and T and A were more frequent than A and C in the second and third SNP, respectively.

**Figure 5.**
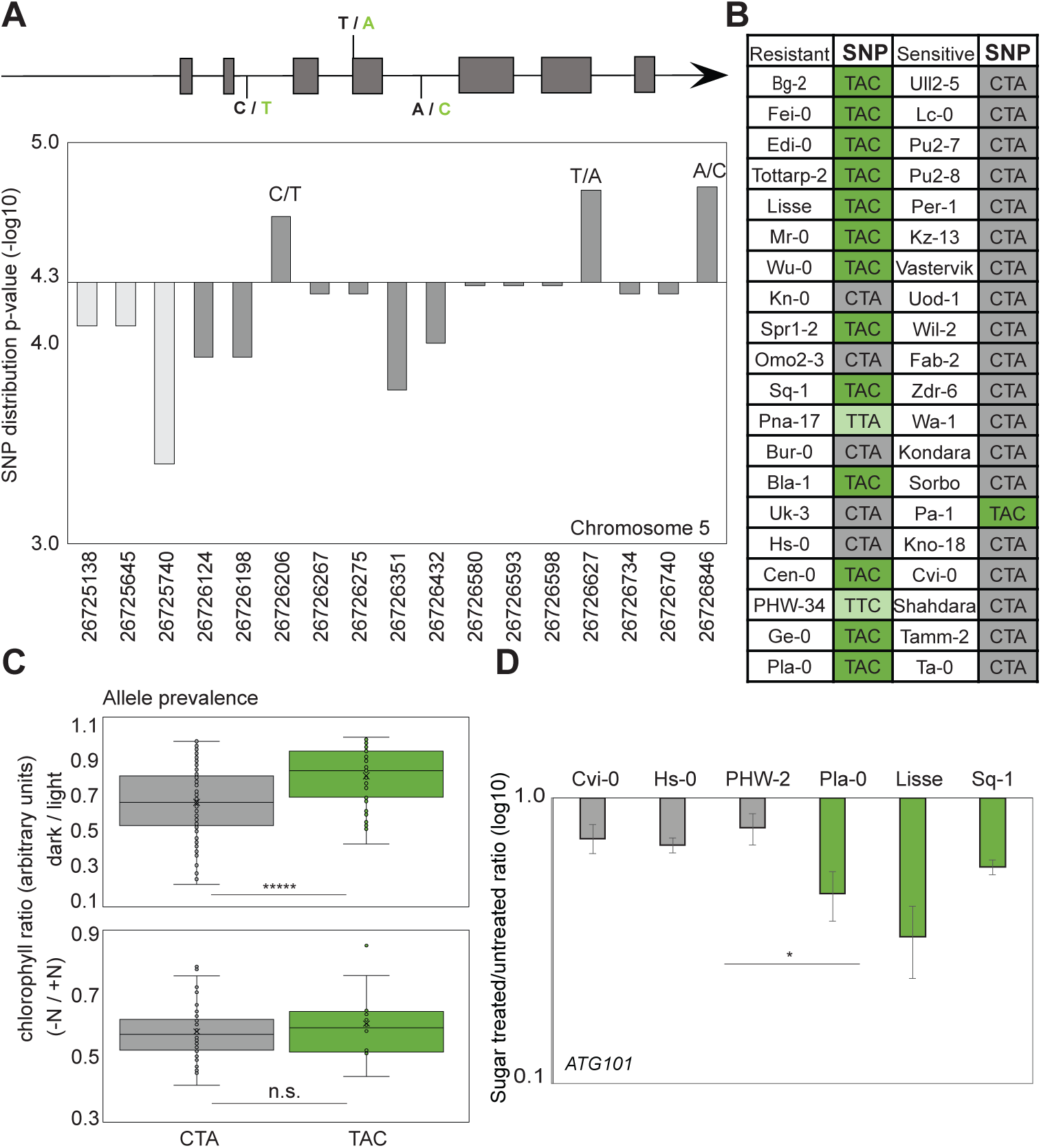
SNPs found in *ATG101* genomic region (promoter-1500 bp upstream and genomic sequence) in a range of 181 *A. thaliana* accessions. A. Upper panel: schematic representation of *ATG101* gene together with the position of the three SNPs showing the highest significance in relation to the dark / light chlorophyll ratio; lower panel: graphic representation of the SNPs within *ATG101* gene together with their p-value (log10); only SNPs with minor allele frequency (MAF) of 15 or more were considered. B. Table depicting the occurrence of the two main allele combinations TAC and CTA of the three SNPs with the highest significance in relation to the perceived resistance/ sensitivity to prolonged fixed carbon starvation; grey represent the allele associated with sensitivity while dark green the one associated with resistance; light green is used to indicate the accessions with intermediate combination of SNPs. C. Upper panel: Distribution of the dark/light chlorophyll phenotype of the accessions between the allele combinations (TAC and CTA containing 39 and 138 accessions, respectively; p = 5.187×10^-6^); lower panel: (-N/+N) chlorophyll ratio distribution between the two main allele combination groups (differences between the groups are not significant (n.s.; p = 0.44; CTA group contains 38 accessions while TAC group contains 10). D. The comparisons between the SNPs and allele combination groups were performed using two-tail t-test and assuming unequal variance and the differences were found to be statistically significant for p = 0.046.

Two main allele combinations, CTA and TAC, accounted for more than 90% of the accessions (Supplementary Table S1). The CTA allele was strongly associated with lower chlorophyll ratios and was more frequent in CS^S^ accessions while TAC was more frequent in CS^R^ (Figure 5B) and the chlorophyll ratio distribution between the two groups presented significant differences (p = 5.18×^-6^; Figure 5C upper panel). To determine the specificity of the observed response to darkness, we looked at the chlorophyll ratios after nitrogen deficiency and divided them in two groups based on the same allele combinations (CTA and TAC; Figure 5C lower panel). Out of the 51 accessions used in the (-N) chlorophyll experiment, 48 were in the CTA and TAC groups (38 and 10, respectively). No significant differences were found between the two groups indicating that the association between this particular SNP combination and chlorophyll retention might be specific to carbon starvation (Figure 5C lower panel).

To investigate whether these SNPs contribute to carbon availability response, we then probed the response of six accessions to sucrose treatment; three harboring the CTA allele combination (Cvi-0, Hs-0 and PHW-2) and three with the TAC version (Pla-0, Lisse and Sq-1). After a 4-hour treatment of 1-week old seedlings from each accession with 4 % sucrose the *ATG101* gene expression in the seedlings was suppressed and a statistically significant difference between the two groups was found. Therefore, the response to sugar, at least in the investigated accessions, was dependent on the SNP combination. It is worth noting, that the SNP in the fourth intron disrupts a sugar-related cis-element (CATCC) and might therefore affect sugar responsiveness although there are multiple other elements remaining intact.

### Diverse parameters contribute to chlorophyll ratio variation

To assess the impact of several parameters, associated with the accessions, on the perception of carbon starvation and the chlorophyll ratio, we performed a multiple regression analysis (Table 1). For the purpose, flowering time, yearly rain in mm, hours of sunshine, latitude at the place of origin, and number of SNPs (0, 1, 2 or 3) were included. Flowering time had the following categories: early, early/intermediate, intermediate, intermediate/late, late, late/requiring vernalization and requiring vernalization which were given 1, 1.5, 2, 2.5, 3, 3.5 and 4, respectively (based on information at the website of the European Arabidopsis Stock Centre NASC corroborated during propagation). Flowering time and SNPs appeared to be significant factors for the carbon starvation-induced chlorophyll phenotype with p-values of 0.02818152 and 2.8724×10^-5^, respectively. Their contribution however, as deduced by the calculated coefficient, was relatively low (Table 1). It should be noted, that the coefficient of the intercept was the highest (0.69405; p = 4.4652×10^-5^) which is not unexpected since many environmental, physiological and developmental parameters pertaining to the accessions were not included in this analysis due to being inaccessible. The overall significance of the analysis was estimated to be p = 0.0003215.

**Table 1.** Multiple regression analysis of factors with potential influence on the carbon starvation chlorophyll phenotype.

| Regression Statistics |  |  |  |  |  |
| --- | --- | --- | --- | --- | --- |
| Multiple R | 0,350346419 |  |  |  |  |
| R Square | 0,122742613 |  |  |  |  |
| Adjusted R Square | 0,097678117 |  |  |  |  |
| Standard Error | 0,171268043 |  |  |  |  |
| Observations | 181 |  |  |  |  |
| ANOVA |  |  |  |  |  |
|  | *df* | *SS* | *MS* | *F* | *Significance F* |
| Regression | 5 | 0,718222574 | 0,14364451 | 4,89707073 | **0,000321501** |
| Residual | 175 | 5,133229944 | 0,02933274 |  |  |
| Total | 180 | 5,851452518 |  |  |  |

|  | *Coefficients* | *Standard Error* | *t Stat* | *P-value* | *Lower 95%* | *Upper 95%* | *Lower 95.0%* | *Upper 95.0%* |
| --- | --- | --- | --- | --- | --- | --- | --- | --- |
| Intercept | **0,694050824** | 0,165750343 | 4,18732663 | **4,4652E-05** | 0,366923882 | 1,02117777 | 0,36692388 | 1,02117777 |
| flowering | **0,035704878** | 0,016133163 | 2,21313561 | **0,02818152** | 0,003864266 | 0,06754549 | 0,00386427 | 0,06754549 |
| yearly rain | 3,11351E-05 | 3,65886E-05 | 0,85095096 | 0,39595944 | -4,1077E-05 | 0,00010335 | -4,1077E-05 | 0,00010335 |
| hrs sunshine | -4,05417E-05 | 3,60246E-05 | -1,125391 | 0,26196447 | -0,00011164 | 3,0557E-05 | -0,00011164 | 3,0557E-05 |
| latitude | -0,000943527 | 0,002146337 | -0,4395989 | 0,66077002 | -0,00517956 | 0,00329251 | -0,00517956 | 0,00329251 |
| SNP | **0,045879592** | 0,010678253 | 4,29654468 | **2,8724E-05** | 0,024804859 | 0,06695433 | 0,02480486 | 0,06695433 |

We then looked at the *ATG101* promoters and introns in other plant species for cis-elements by focusing on the same ten elements as earlier (Supplementary Figure S4). All of the examined *ATG101* orthologues in different plant species have seven or more of the ten investigated elements indicating that sugar-mediated regulation of *ATG101* gene is conserved in the plant lineage.

### *ATG101* SNP variants complement *atg101* to a different extent

To establish the impact of the SNPs on the regulation of *ATG101* expression and autophagy induction, we decided to investigate further the correlative links that were established by implementing *in planta* changes in the gene sequence. In order to test the involvement of the three SNPs most significantly correlating with the resistance/sensitivity phenotype of the accessions in the regulation and function of *ATG101* gene, we created three different complementation constructs (Figure 6A). The first one contained *ATG101* promoter (-1500 bp) and genomic sequence At5g66930.2 (*ATG101* gene), version of Col-0 which has the TAC allele combination (C lines). The second construct was obtained by mutating the first to match the two intronic SNPs (T to C and C to A) via side-directed mutagenesis (Cm lines). For the third construct, the Kz-9 accession version of *ATG101* was used, complete with a promoter from the same accession, giving rise to the Ci lines. Constructs were introduced into the *atg101* mutant (Wiscseq_DsLox337F01) (Lee et al., 2023) through floral dip transformation and T2 lines, confirmed to have normal development and to be expressing the gene, were used for all the analyses (Figure 6).

**Figure 6.**
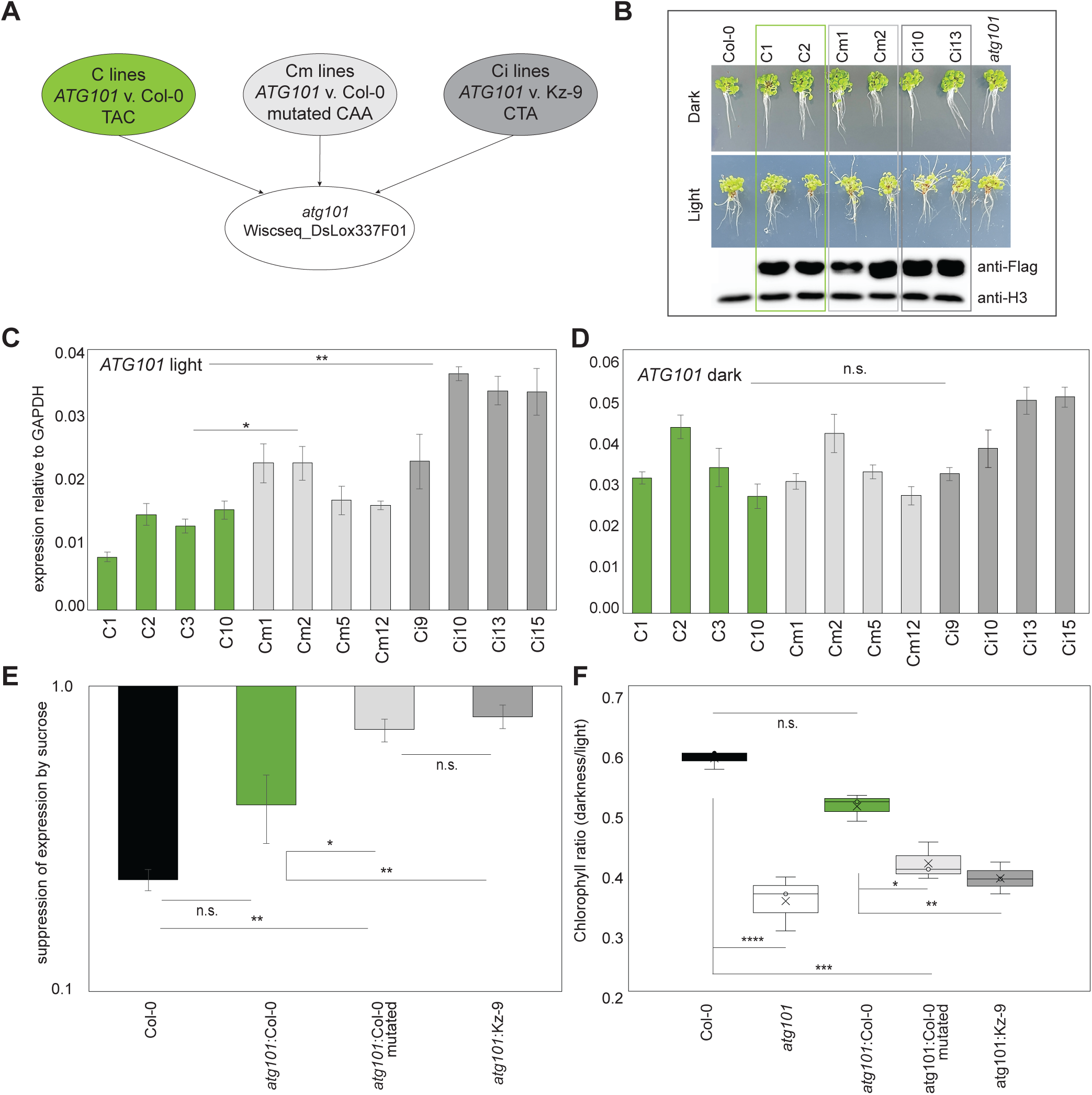
Complementation of the *atg101* mutant phenotype with constructs containing different SNP alleles leads to different results. A. *atg101* mutant complementation: scheme explaining the used constructs and obtained lines; from left to right construct containing either the Col-0 version of the gene (C; TAC), the mutated Col-0 version (Cm; CAA) or a Kz-9 version of the promoter (-1500 bp) and gene (Ci; CTA). B. Upper panel: 7-day old seedlings of WT Col-0, different complementing lines (two of C, Cm and Ci complementing lines each) and *atg101* mutant; middle panel: 1-week dark treated seedlings from the same plant lines as above; lower panel: Western blot analysis determining the expression levels of ATG101 protein in light grown seedlings (3 x Flag tagged); histone H3 antibody was used as a loading control. C. *ATG101* gene expression under control (light) conditions in independent C, Cm and Ci complementing lines. D. *ATG101* gene expression in complementing lines (same as in C) after a 24 h dark treatment. E. Differences in the responsiveness of the *ATG101* gene expression to sucrose treatment in Col-0 and *atg101:ATG101* complementing lines. F. Differences in the complementation of the autophagy defect related chlorophyll loss phenotype of *atg101* mutants done with different variants of *ATG101* gene. Data in C, D and E are derived from two independent experiments involving each of the independent lines used. Data in F are derived from three independent experiments for each of the individual lines involved. Statistical analysis was performed using ANOVA, Tukey test including a pairwise comparison test to probe the differences between the groups investigated. Statistically significant differences between the means are indicated with asterisks as follows: * for p<0.05, ** for p<0.005, *** for p<0.0005 and **** for p<0.00005. Error bars in C, D and E indicate standard error calculated for biological replicates while in F they indicate positive and negative variation..

The expression of *ATG101* gene was analyzed in multiple lines resulting from the transformation of each construct. For further experiments were chosen four lines of each genotype which exhibited similar levels of the gene after a dark treatment (a feature of all the examined accessions; level of expression, not induction). Normal seedling development and expression of the ATG101 protein were also confirmed (Figure 6B). All 12 lines were examined for the expression of *ATG101* in control seedlings and after dark treatment (Figure 6C and D). The transcriptional levels of *ATG101* in Cm and Ci lines under normal conditions were higher than in C lines. This result is in line with our observation that accessions containing TAC allele combination of SNPs tend to have lower levels of *ATG101* in control seedlings than accessions containing the CTA allele. Statistically significant differences were shown for the C lines compared to both Cm and Ci by applying ANOVA Tukey test followed by a pairwise test (* p < 0.05 for Cm – C and ** p < 0.005 for Ci – C, respectively; Figure 6C). No significant differences among the three groups in the *ATG101* expression induced by darkness were found (Figure 6D).

Next, we investigated the effect of 2% sucrose on the three different sets of complementing lines (Figure 6E). The strongest suppression of *ATG101* after 2 h of treatment was observed for the C lines. Col-0 seedlings exhibited somewhat stronger inhibition but the differences with C lines were not significant. Neither the Cm, nor the Ci lines showed a comparable suppression of *ATG101* gene expression (levels for both sets of lines had statistically significant differences from WT and the C lines; * p < 0.05 and ** p < 0.005 respectively; Figure 6E). Additionally, we tested the level of complementation achieved by the introduction of the three constructs by subjecting T2 seedlings from all 12 independent lines to a prolonged dark treatment and analyzing the chlorophyll. The C lines complemented the *atg101* mutant and their chlorophyll ratio levels showed no significant differences from the wild type (Figure 6F). The chlorophyll content in the complementing lines after a dark treatment corresponded to the observed phenotypic changes (Figure 6B upper panel). The Cm and Ci lines, on the other hand, although showing increased chlorophyll levels compared to the *atg101* mutant, still had significantly lower ratios than WT (Figure 6F).

Neither the responsiveness to sugar, nor the chlorophyll phenotype after a prolonged dark treatment were restored to WT levels in *atg101* seedlings with a mutated Col-0- derived *ATG101* construct (CAA allele combination) or a Kz-9 derived one (CTA allele combination). Only the construct containing the TAC allele combination, associating strongly with carbon starvation resistant natural accessions, was able to restore both parameters to WT levels. Thus, the presence of SNPs in the *ATG101* genomic region appear to play a significant role for the regulation of *ATG101* gene and by extension autophagy induction and regulation in the context of natural variants.

Based on the conducted study and focusing largely on the interplay between *ATG101* expression level, sugar responsiveness, SNPs, and their potential effect on autophagy initiation, we propose the following model (Figure 7). In CS^R^ accessions *ATG101* expression in light grown seedlings is low, thus maintaining low baseline level of autophagy due to reduced initiation complex assembly. CS^S^ accessions have overall higher control *ATG101* expression which would determine more elevated baseline autophagy. After a dark treatment both types of accessions induced *ATG101* gene expression. The induction appeared to be stronger in CS^R^ accessions which contributes to higher autophagy efficiency making them more resistant to prolonged darkness. The differential induction of *ATG101* extends also to the TOR kinase which exhibits stronger suppression in CS^R^ lines. The extent of TOR inhibition has been shown to impact autophagy initiation, with stronger TOR suppression correlating to stronger autophagic responses (Dong et al., 2022). Complementation of the *atg101* mutant phenotype appears to be more successful when the TAC allele was used thus once again indicating the role of the SNPs in *ATG101* locus in determining prolonged darkness sensitivity.

**Figure 7.**
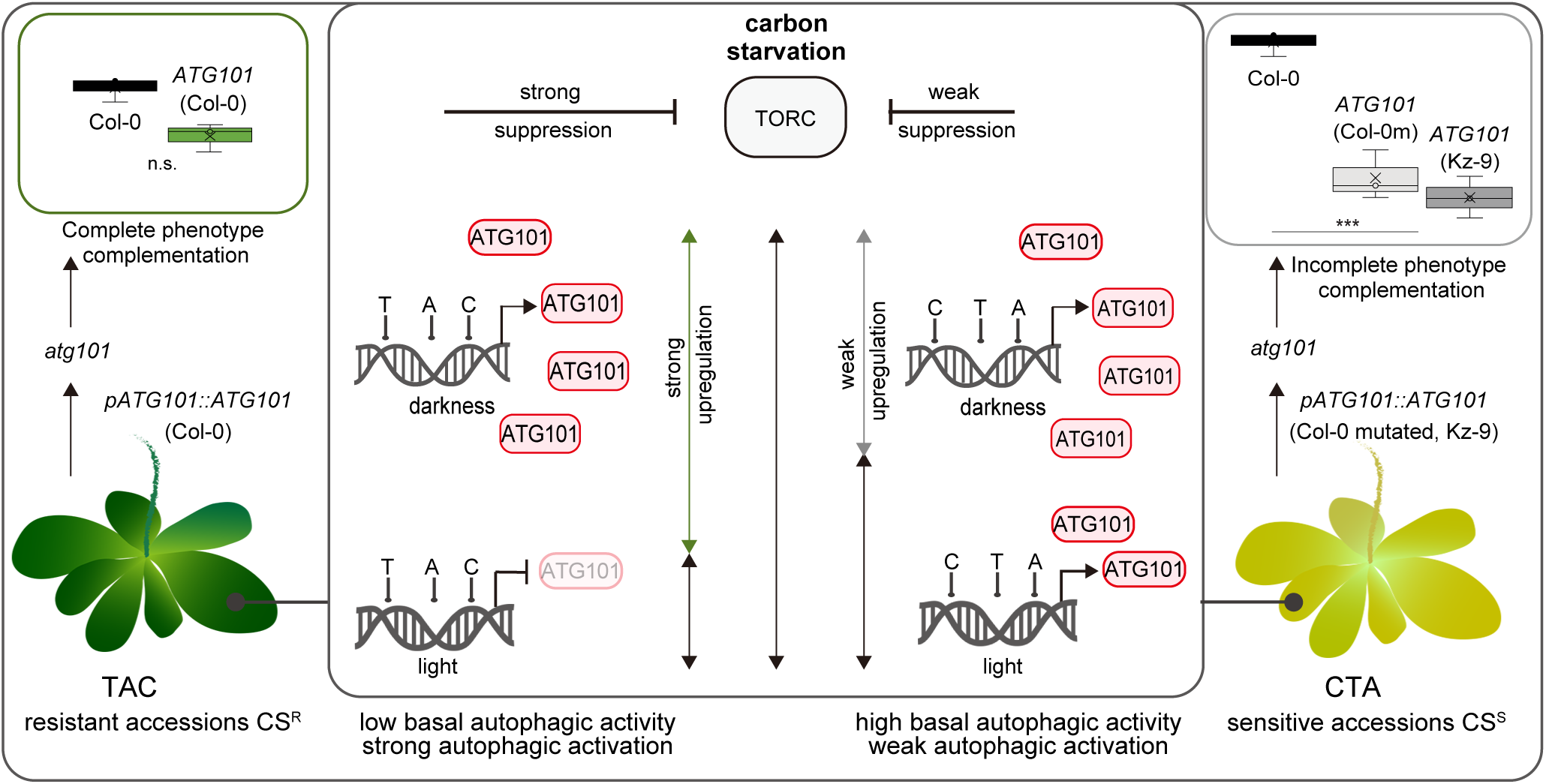
Proposed model. In CS^R^ accessions *ATG101* expression in light grown seedlings is low, thus maintaining low baseline level of autophagy due to the lack of initiation complex assembly. CS^S^ accessions have overall higher control *ATG101* expression which would determine more elevated baseline autophagy. After a dark treatment both type of accessions induced *ATG101* gene expression. The induction appeared to be stronger in CS^R^ accessions which contributes to higher autophagy efficiency making them better in withstanding prolonged darkness. The differential induction behavior pertaining to *ATG101* extends further upstream to the TOR kinase which exhibits stronger suppression in CS^R^ lines. The differential regulation of the *ATG101* gene and the differences in functionality are illustrated by the different outcomes resulting from mutant complementation with different variants of the gene. The Col-0 (TAC) version is able to fully complement the mutant phenotype while the mutated Col-0 or the Kz-9 versions brought about only partial complementation of the *atg101* mutant.

Our results demonstrate that *ATG101* is suppressed by various sugars, suggesting that under optimal photosynthetic productivity, it is not transcribed in large mounts. Carbon starvation, but not nitrogen deprivation, triggers its strong induction. Resistance to carbon starvation does not translate into an ability to withstand prolonged nitrogen deficit and *vice versa*. Two main SNP allelic combinations in the *ATG101* locus were shown to contribute to the differential regulation of *ATG101* and the reactivity of Arabidopsis accessions to carbon starvation.

## Discussion

### Sugar-related modulation of *ATG* genes and natural variation of autophagy

Autophagic degradation is a pathway universally present in eukaryotes which is involved in the elimination of unwanted molecules and organelles (Liu and Bassham, 2012; Yang and Bassham, 2015; Havé et al., 2017; Masclaux-Daubresse et al., 2017; Marshall and Vierstra, 2018) and thus it is important for the maintenance of cellular homeostasis as well as plant growth, development, metabolic processes and environmental responses (Thompson et al., 2005; Li et al., 2015; Havé et al., 2017; McLoughlin et al., 2020).

Plants are particularly vulnerable to changing environmental conditions due to their sessile nature making adaptations, such as adjustments in nutrient recycling, crucial for survival. *A. thaliana* natural variants inhabit a wide variety of conditions and not only survive but are able to thrive, while staying within the constraints of the species definition. It is conceivable, that accessions have acquired numerous adaptations, tailored to the specific conditions at their place of origin. In this study we have focused on the variation of carbon starvation-induced autophagy and its regulation through *ATG101*.

Autophagy in plants has been demonstrated to serve as an energy provider for growth during the dark period and it is downregulated by either switching the plants to continuous light or providing them with exogenous sugar (Izumi et al., 2013). It has long been established that sugar not only serves as a source of energy but also as a signaling molecule (Sheen et al., 1999; Li and Sheen, 2016). Some proteins are able to act as sugar sensors such as, for instance, the hexokinase 1 (HXK1) which has preference towards glucose (Kawai et al., 2005). Soluble sugars, like sucrose and different hexoses, can modulate gene expression in a more or a less specific manner (Price et al., 2004; Wind et al., 2010; Li and Sheen, 2016) by interacting with cis-regulatory elements in the promoters of the respective genes in reaction to sugar starvation or supply (summarized in (Sakr et al., 2018)). Approximately 10 % of the Arabidopsis genome consists of sugar-responsive genes (Price et al., 2004).

The TOR kinase, a negative regulator of autophagy upstream of the initiation complex (Liu and Bassham, 2010), is also dependent on sugars such as glucose and sucrose (Huang et al., 2019; Li and Zhao, 2024). In this study, we demonstrated that ATG1 – ATG13 complex encoding genes are modulated at the transcriptional level by various sugars. We have shown that Arabidopsis accessions exhibit clear differences in terms of darkness-induced autophagy. Senescence, another prolonged darkness-triggered processes, also exhibits natural variation which appears to be related to several amino acids and enzymes involved in their metabolism (Zhu et al., 2022). Another trend we observed, namely the diversified but different response of the accessions to nitrogen starvation and the lack of major induction of *ATG101* gene expression by N-deficit, indicated a certain level of specificity of the responses to darkness.

### *ATG101* sequence variation – sugar interplay regulates darkness-mediated autophagy induction

The potential of ATG101 to regulate autophagy initiation has been recognized in mammals as it forms a HORMA heterodimer with ATG13 and the complex acts as a hub for interaction with multiple other factors (Qi et al., 2015). In Arabidopsis, crossing *atg101* to glycolysis mutants was able to suppress the spontaneous induction of autophagy in the latter, establishing simultaneous connections for ATG101 with sugar metabolism and autophagy regulation (Lee et al., 2023). Therefore, the involvement of ATG101 in determining natural variation of sensitivity to prolonged darkness investigated here is another aspect of ATG101 regulatory role which has not been studied so far.

It should be noted, that while the two investigated SNP alleles, TAC and CTA, correlate strongly with resistant and sensitive accessions, there are exceptions, particularly in the resistant part where multiple accessions carry the CTA combination. Other factors and/or other features of *ATG101*, such as indels and structural variations (defined as ˂50 bp and ˃50 bp nucleotide changes, respectively), could contribute to these differences. One version of such indels is however present in the Ci complementing lines (Kz-9 version of the gene). While no improvement or exacerbation of the phenotype resulted from using this version of the gene in comparison to the effect seen in Cm (mutated Col-0 version), neither outcome can be excluded in other accessions where structural variations are present. The disruption of one of the sugar-related cis-regulatory elements in the fourth intron by a SNP has also potential to influence not only the responsiveness to sugar but the overall functionality of *ATG101*. The significance of polymorphisms in cis-elements for the expression of the respective genes has already been demonstrated (Wang et al., 2021). It should also be emphasized that in the current study we only investigate one of the possible mechanisms involved in determining sensitivity to carbon starvation.

### Factors of potential consequence in determining sensitivity to carbon starvation

Multiple endogenous and exogenous factors are likely to determine carbon starvation-induced autophagy efficiency of the Arabidopsis accessions. The accessions used in this study have high variability in terms of flowering time with some of them needing long periods of cold treatment (vernalization) to be able to flower. The flowering time was therefore included in a multiple regression analysis where it proved relevant to the chlorophyll ratio variation and, by extension, the carbon starvation sensitivity. The *SOC1* transcription factor which is involved in transition to flowering has already been linked to autophagy as it suppresses *ATG* gene expression (Li et al., 2022). Additionally, subjecting seedlings to darkness just before flowering renders them unable to provide the necessary energy and hormonal flux for the transition to reproductive development (Sheikh et al., 2024), or alters the expression of light-responsive genes essential for flowering (Armarego-Marriott et al., 2020). Therefore, the correlation with flowering time in our study is unsurprising. The other significant factor determining carbon starvation sensitivity was the SNP allele combination present in the individual accessions (focusing on the three SNPs most significantly associated with the chlorophyll phenotype). The importance of the SNPs for the overall manifestation of the phenotype of the accessions after carbon starvation was also experimentally investigated and has already been discussed.

Finally, by looking at the *ATG101* promoter and gene in other organisms, we noticed that not only the Arabidopsis locus contains promoter bound sugar regulatory elements but also *ATG101* loci from other clades. Practically, all of the examined *ATG101s* have most types of the cis-elements we investigated. The observed high level of conservation of the sugar related cis-regulatory elements might be an indicator of their significance for the gene regulation. Further research, combining genetic, evolutionary, developmental and physiological approaches, would certainly improve the understanding of natural variation of autophagy.

## Materials and methods

### Plant growth and treatment

*Arabidopsis thaliana* seeds from 181 different accessions, including Columbia-0 (Col-0), *atg101* mutant (Wiscseq_DsLox337F01), and complementing lines in the same background were sterilized using three consecutive rinses with ethanol (two times 70% and once 100%) on spin columns followed each time by a brief centrifugation at 14000 rpm. The seeds were left to dry in sterile conditions and sown on ½ MS square plates containing 0.8 % agar. Plates were stratified for 48 h at 4 °C and placed vertically or horizontally in a growth chamber Panasonic. The plants were grown at short-day (8 h day, 16 h night) settings with 120 μmol PPFD, 22 °C day / 18 °C night temperature for both carbon and nitrogen starvation, 7 and 21 days, respectively. For a dark treatment, plants were later transferred for 24 h or 7 days in darkness. Seedlings were usually put in darkness at 10 am or 2 h after the beginning of the day period. Control plants were harvested at the time of the transfer, typically within 30 min. Plants for nitrogen deficiency treatment were grown for one week on normal ½ MS medium and then transferred for 48 h or 14 days at ½ MS medium without nitrogen. The samples for gene expression and protein analysis were immediately frozen in liquid nitrogen and later analyzed while the ones for chlorophyll extraction were gently dried with paper towels, weighed and processed further. All plant samples taken from dark-treated plants were processed according to their purpose.

### Sucrose, glucose, galactose, arabinose, and mannitol treatment

Wild-type (WT) Col-0 were plated on ½ MS medium without sugar and after one week the seedlings were transferred to ½ MS medium containing 2 % sugar or 25 mM mannitol. Control seedlings were transferred to medium without any supplement.

Samples for monitoring gene expression were taken from control and treated seedling at 0 h, 1 h and 2 h after the treatment. For the sugar response related to the SNP alleles, selected accessions were grown for 7 days on ½ MS agarose medium without sugar. The seedlings were then transferred to ½ MS medium supplemented with 4% sucrose, or to medium without sugar. Samples were taken 4 h after the transfer from both control and sugar-treated seedlings followed by gene expression analysis.

### Chlorophyll quantification

Control, 7-day dark-treated, and 14-day no nitrogen-treated Arabidopsis seedlings were put after weighing in a 96-deepwell polypropylene plate prefilled with 1 ml dimethyl formamide (DMF) in each well. The optimization of the chlorophyll extraction allowed processing of a large number of samples simultaneously. As soon as all the seedlings were added, the plate was sealed with a transparent sticky film, wrapped in aluminum foil and left at 4 °C for 48 h, shaking at 120 rpm for the first 24 h. In order to quantify the chlorophyll content,150 μl of the extract was transferred to a standard 96-well polypropylene plate and the absorption was determined at 664 nm and 647 nm using a microplate reader Tecan M-Plex. For calculating the chlorophyll content of the samples, the following formula was used:

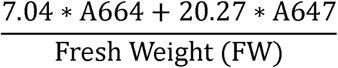

After application of the formula, control and after-treatment chlorophyll values were used to calculate the darkness/light and the (-nitrogen)/(+nitrogen) chlorophyll ratios.

### RNA extraction, reverse transcription, QRT-PCR

Samples frozen in liquid nitrogen were ground using 1.2 mm ∅glass beads and a tissue lyser Qiagen. The RNA Plant and Algae extraction kit of Macherey-Nagel was used for further processing of the samples in accordance with the manufacturer’s instructions. RNA was reverse transcribed using MMulv reverse transcriptase (Promega) according to the enclosed instructions. After the reaction was completed, the resulting cDNA was used for QRT-PCR analysis. *GAPDH* (glyceraldehyde phosphate dehydrogenase) and *YLS8* (yellow leaf specific 8) were employed simultaneously as reference genes in the QRT-PCR analysis. Calculations of the QRT-PCR results were performed using the ΔΔCt method. For each gene and treatment variant at least two independent experiments were performed. The typical number of technical replicates in every experiment was three. Where applicable additional normalizations were used (*i.e.,* the control condition or a specific time point or concentration) as a reference. All used primer sequences, together with the corresponding genes and numbers, are listed in Supplementary Table S2.

### Monitoring TOR kinase activity

Arabidopsis seeds were sown in 6 well plates with 1 ml of ½ MS medium and grown for 7 days under short day conditions (the liquid medium was replaced on days 4 and 6). For the dark-treated samples, the seedlings were collected just before the end of the night period on day 7, and for the light control samples, the seedlings were exposed to light for 5 h on day 7 and collected. The collected seedlings were immediately frozen in liquid nitrogen, ground with 2x Laemmli sample buffer, and denatured for 4 min at 95 °C. The samples were centrifuged at 15,000 × *g* for 2 min, and the supernatant was subjected to Western blot with the anti-RPS6A-P240 antibody (Agrisera: AS19 4302) and anti-RPS6A (Agrisera: AS19 4292).

### WB analysis of atg101 complementing lines

For ATG101 – 3xFlag tag fusion protein samples were taken at the same time as the ones for control gene expression and chlorophyll, frozen in liquid nitrogen and extracted with a sample buffer containing 50 mM Tris-HCl pH 7.5, 150 mM NaCl, 0.5 % Triton X-100 and protease inhibitor cocktail (Roche) and centrifuged for 10 min at 15 000 x *g*. Protein was quantified and denatured by adding the appropriate amount of 6 x Laemmli buffer and boiling the sample at 95 °C for 3 min. To probe ATG101 protein level in the complementing lines, anti-Flag (F 1804; Sigma) antibody was used while the anti-H3 antibody from Agrisera (AS10 710) was used as loading control.

### Cloning and transformation of complementation constructs, and analysis of the resulting T2 stable plant lines

For the complementation of the *atg101* knockout mutant three separate constructs were used. Golden Gate cloning method was used for the cloning. The *ATG101* promoter and gene were amplified using separate sets of primers. 3 x Flag – tag and an OLE (oleosin) promoter-driven RFP selection marker for the seeds were also added to the assembly. The promoter and gene were amplified from Col-0 and Kz-9 accessions. The fragments were inserted into a LIIalphaF1-2 backbone vector as described in (Binder et al., 2014). Subsequently, the Col-0 version of the construct was mutated in the positions of the two intron SNPs through site-directed mutagenesis using two overlapping primers for each mutation to perform a PCR followed by a digestion of the non-mutated template with the help of DpnII restriction endonuclease. Primers are listed in Supplementary Table S2. The three constructs were amplified in *Escherichia coli*, controlled for correct assembly and introduced in the *atg101* mutant (Wiscseq_DsLox337F01) via Agrobacterium-mediated transformation. Seeds positive for the RFP marker were sown and the resulting seedlings were tested for expression of the *ATG101* gene and protein. Multiple independent lines for each construct were used for further experiments in the T2 generation.

### Statistical analysis

Descriptive statistics were used to determine average and standard error values, t-test and ANOVA analysis, usually coupled with a subsequent pairwise comparison, were employed to assess the significance of the differences between accessions and treatment groups, multiple regression was performed to evaluate the effect of multiple dependent variables on an independent one and t-test were carried out to compare whole groups, respectively. Box-plot analysis with and without outliers was used to visualize the entire range of variation of accessions and treatment groups. All statistics was performed using the Xlrealstats add-in of MS Excel. Error bars in bar graphs represent standard error and are always calculated for biological replicates, in Box-plots error bars represent positive and negative variation. For all t-tests unequal variance between the sample groups was assumed. Pearson correlation analysis was employed to analyze the possible relationship between chlorophyll ratios after dark treatment and nitrogen deprivation of the same sampling groups.

### Analysis of promoters and introns of autophagy related genes for the presence of cis-regulatory elements

Promoter (1500 bp upstream of the transcription start site) and genomic sequences of *ATG101*, *ATG1* and *ATG13* homologues, and *ATG11* were taken from Phytozome.org. All sequences were analyzed for the presence of regulatory elements manually and using https://www.dna.affrc.go.jp/PLACE/ and manually. Elements related to sugar signaling were indicated with a kind and a numerical value for A. thaliana and only with presence/absence for all the other species. In the cases where multiple homologues were present in a particular species only one was chosen at random.

## Acknowledgements

We thank the Alexander von Humboldt Foundation and the Zukunftskolleg of the University of Konstanz for financial support for Svetlana Boycheva Woltering. Joost Woltering is acknowledged for commenting on the manuscript. Kentaro Shimizu and Rie Shimizu-Inatsugi are acknowledged for providing the natural accessions of *Arabidopsis thaliana*

## Author contributions

SBW and EI conceived the study. SBW performed and analyzed most of the experiments. MT and TB performed and analyzed some of the experiments. SBW and EI wrote and finalized the manuscript. All authors have read and agreed with the final version of the manuscript.

