## Supplementary material for "Sugar-mediated control of *ATG101* modulates carbon deficiency-induced autophagy in Arabidopsis"

**Figure S1.** Sugar related cis-regulatory elements in *ATG8* homologues and response to sugar.

**A.** Scheme of the presence/absence of ten different sugar-related elements in promoters or introns of three different ATG8 homologues.

**B.** Effect of treatment with 1 % sugar on the expression of *ATG8c, ATG8e* and *ATG8f* genes. The scale is logarithmic and the results are mean values of two independent experiments containing one biological and three technical replicates each. Error bars represent standard error as calculated for the biological replicates.

**Figure S2.** Remaining 131 dark-treated/control chlorophyll ratios of the *A. thaliana* accessions.

**Figure S3.** TOR kinase activity.

**A.** Average TOR kinase activity for seedlings of eight (4 CSR and 4 CSS) light-grown *A. thaliana* accessions.

**B.** Average TOR kinase activity for seedlings of eight dark-treated A. thaliana accessions. Data are mean values of 3 independent experiments. Error bars on the box-plots represent positive and negative variations. No significant differences could be found between the resistant and sensitive groups.

**Figure S4.** Conservation of sugar-related cis-regulatory elements in promoters and introns of different plants species.

**A**

| | CGACG | TATCCA | TTATCC | TATCCAY | CATCC | GGATA | ATATCT | GATAA | GATTA | ACGTG |
|---|---|---|---|---|---|---|---|---|---|---|
| *ATG8c* | | | | | | | | 7 | 6 | |
| *ATG8e* | | 1 | 2 | 1 | 1 | 2 | 3 | 9 | 5 | 6 |
| *ATG8f* | | 2 | 2 | 2 | 1 | 8 | 4 | 12 | 9 | 5 |

**B**

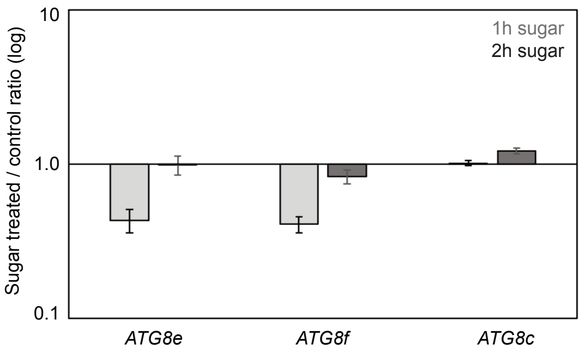


Figure S1. Sugar related cis-regulatory elements in *ATG8* homologues and response to sugar. A. Scheme of the presence/absence of ten different sugar-related elements in promoters or introns of three different ATG8 homologues. B. Effect of treatment with 1 % sugar on the expression of *ATG8c, ATG8e* and *ATG8f* genes. The scale is logarithmic and the results are mean values of two independent experiments containing one biological and three technical replicates each. Error bars represent standard error as calculated for the biological replicates.

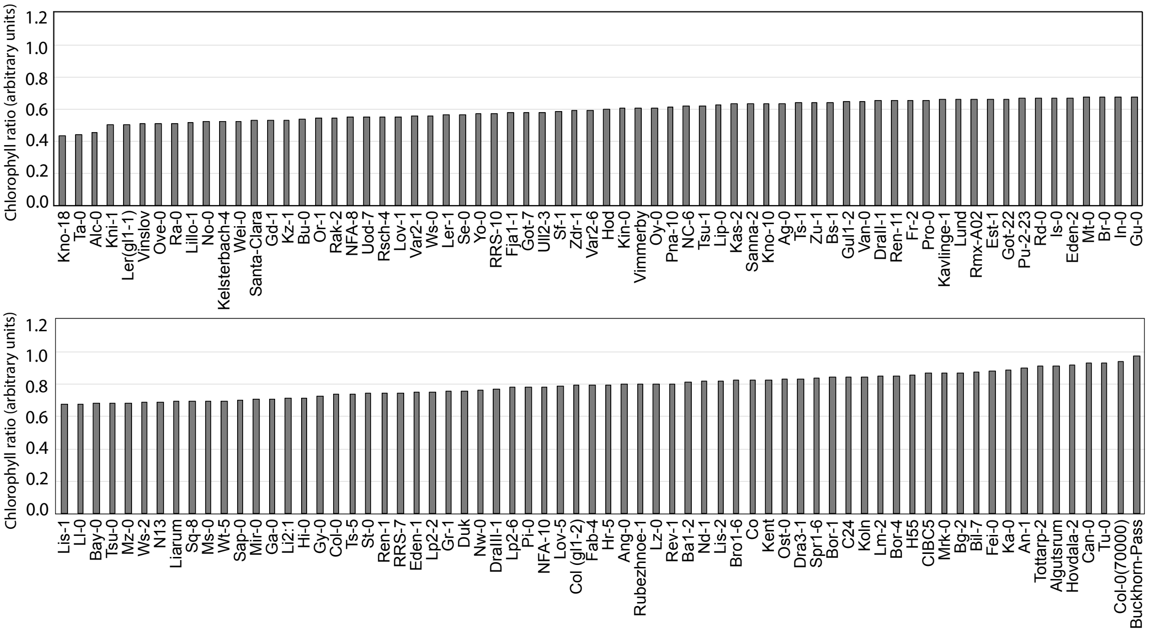


Figure S2. Remaining 131 dark-treated / control chlorophyll ratios of the *Arabidopsis thaliana* accessions.

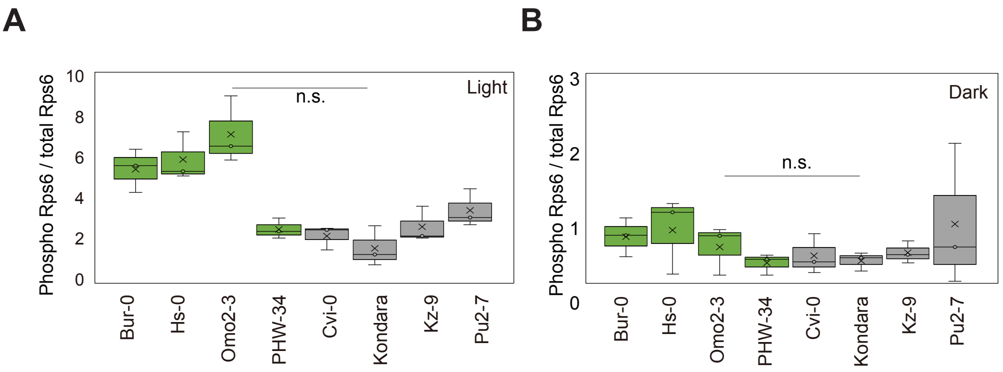


Figure S3. TOR kinase activity. A. Average TOR kinase activity for seedlings of eight (4 $CS^R$ and 4 $CS^S$) light-grown *A. thaliana* accessions. B. Average TOR kinase activity for seedlings of eight dark-treated *A. thaliana* accessions. Data are mean values of 3 independent experiments. Error bars on the box-plots represent positive and negative variations. No significant differences could be found between the resistant and sensitive groups.

| Conservation of *cis* regulatory elements in *ATG101* gene promoters and introns | | | | | | | | | | |
|---|---|---|---|---|---|---|---|---|---|---|
| + | | | | | + | | | | | CGACG |
| + | + | + | + | + | + | + | + | + | + | TATCCA |
| | + | + | | | + | + | | | + | TTATCC |
| + | + | + | + | + | + | + | + | + | + | TATCCAY |
| + | + | + | | + | + | + | + | + | + | CATCC |
| | + | + | + | + | + | + | + | + | + | GATAA |
| + | + | + | + | + | + | + | + | + | + | ATATCT |
| + | + | + | + | + | + | + | + | + | + | GGATA |
| + | | + | + | + | + | + | + | + | + | GATTA |
| + | + | + | + | + | + | | + | + | + | ACGTG |

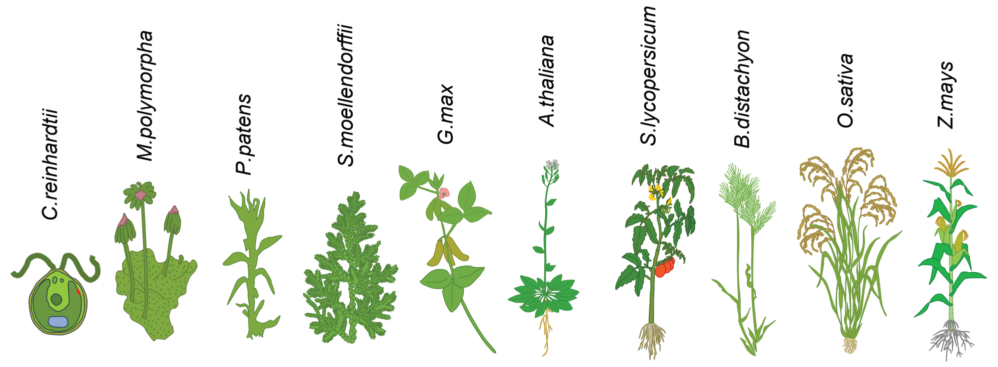


Figure S4. Conservation of sugar-related cis-regulatory elements in promoters and introns of different plants species.

**Supplementary Table 1**

| name | (-C)chl.ratio | flowering | yearly rain | hrs sunshine | latitude | SNP | |
|---|---|---|---|---|---|---|---|
| Ag-0 | 0.63380004 | 3.5 | 1025 | 2000 | 45.2 | 0 | CTA - 0 |
| Alc-0 | 0.456553325 | 3.5 | 423 | 2744 | 40.3 | 0 | TAC - 3 |
| Algutsrum | 0.911640255 | 3 | 400 | 2092 | 56.7 | 0 | |
| An-1 | 0.900293306 | 1 | 825 | 1630 | 51.5 | 0 | |
| Ang-0 | 0.798361444 | 1 | 139 | 1670 | 50.5 | 0 | |
| Ba1-2 | 0.814593923 | 1 | 924 | 2366 | 56.4 | 1 | |
| Bay-0 | 0.680270728 | 2 | 957 | 2251 | 40.2 | 0 | |
| Bg-2 | 0.871690982 | 2 | 999 | 2170 | 47.6 | 3 | |
| Bil-5 | 0.890302398 | 4 | 372 | 2358 | 63.3 | 0 | |
| Bil-7 | 0.874226651 | 4 | 372 | 2358 | 63.3 | 0 | |
| Bla-1 | 0.965750733 | 3 | 720 | 2330 | 41.5 | 2 | |
| Bor-1 | 0.84235981 | 2 | 510 | 1840 | 49.4 | 0 | |
| Bor-4 | 0.850712691 | 2 | 510 | 1840 | 49.4 | 0 | |
| Br-0 | 0.67514061 | 3.5 | 510 | 1840 | 49.2 | 0 | |
| Bro1-6 | 0.824504857 | 3.5 | 325 | 2427 | 56.3 | 3 | |
| Bs-1 | 0.638504072 | 1 | 842 | 1687 | 47.5 | 0 | |
| Bu-0 | 0.535825786 | 1.5 | 1005 | 2166 | 50.5 | 0 | |
| Buckhorn Pas | 0.974218986 | 3.5 | 531 | 3166 | 41.4 | 0 | |
| Bur-0 | 0.965185108 | 2 | 1525 | 3310 | 54.1 | 0 | |
| C24 | 0.842953243 | 2 | 905 | 2795 | 40.2 | 3 | |
| Can-0 | 0.928159616 | 3 | 111 | 2986 | 29.2 | 0 | |
| Cen-0 | 0.97890342 | 1 | 740 | 1690 | 49.2 | 3 | |
| CIBC-17 | 0.922173022 | 2 | 452 | 2199 | 51.4 | 3 | |
| CIBC-5 | 0.866545999 | 2 | 452 | 2199 | 51.4 | 3 | |
| Co | 0.824711856 | 3 | 905 | 2795 | 40.2 | 0 | |
| Col-0 | 0.83633821 | 1.5 | 1120 | 2980 | 38.3 | 3 | |
| Ct-1 | 0.930400062 | 2 | 500 | 3372 | 37.3 | 0 | |
| Cvi-0 | 0.432672629 | 1.5 | 24 | 2925 | 15.1 | 0 | |
| Dra3-1 | 0.829856825 | 3.5 | 1080 | 3299 | 55.8 | 0 | |
| DraII-1 | 0.64996485 | 3 | 522 | 1775 | 49.4 | 0 | |
| DraIII-1 | 0.768729684 | 2 | 522 | 1775 | 49.4 | 0 | |
| Duk | 0.757473163 | 1 | 522 | 1775 | 49.1 | 1 | |
| Eden-1 | 0.749138662 | 4 | 304 | 2582 | 62.9 | 0 | |
| Eden-2 | 0.668819677 | 4 | 304 | 2582 | 62.9 | 0 | |
| Edi-0 | 0.893072397 | 3.5 | 730 | 1450 | 55.9 | 3 | |
| Eds-1 | 0.940534382 | 3 | 721 | 2490 | 62.9 | 0 | |
| Ei-2 | 0.889589866 | 2 | 678 | 1544 | 50.3 | 0 | |
| Est-1 | 0.661450231 | 2 | 687 | 1835 | 58.3 | 0 | |
| Fab-2 | 0.377370922 | 3.5 | 304 | 2582 | 63.0 | 0 | |
| Fab-4 | 0.794188563 | 3.5 | 304 | 2582 | 63.0 | 0 | |
| Fei-0 | 0.883417848 | 2 | 1237 | 2468 | 40.9 | 3 | |
| Fja1-1 | 0.574456198 | 3 | 562 | 2336 | 56.1 | 0 | |
| Fr-2 | 0.650543842 | 1.5 | 600 | 1720 | 50.0 | 0 | |
| Ga-0 | 0.709503969 | 2 | 732 | 2251 | 50.3 | 0 | |
| Gd-1 | 0.528978113 | 1 | 804 | 2247 | 53.5 | 2 | |
| Ge-1 | 0.993656692 | 3 | 925 | 1930 | 46.5 | 3 | |
| Got-22 | 0.662359194 | 3.5 | 804 | 2193 | 51.5 | 0 | |
| Got-7 | 0.576559097 | 3.5 | 804 | 2193 | 51.5 | 0 | |
| Gr-1 | 0.75745039 | 2 | 810 | 2892 | 47.0 | 0 | |
| Gu-0 | 0.677099398 | 2 | 732 | 2251 | 50.3 | 3 | |
| Gul1-2 | 0.649550742 | 3 | 676 | 2302 | 56.5 | 0 | |
| Gy-0 | 0.727207564 | 2 | 720 | 2266 | 49.5 | 3 | |
| H55 | 0.853719044 | 2 | 732 | 2475 | 49.1 | 0 | |
| Hi-0 | 0.716002592 | 1 | 846 | 2314 | 52.5 | 0 | |
| Hod | 0.597304478 | 3.5 | 822 | 2184 | 48.8 | 0 | |
| Hovdala | 0.918190269 | 2 | 821 | 2159 | 56.1 | 0 | |
| HR-5 | 0.794385267 | 2 | 452 | 2199 | 51.4 | 3 | |
| Hs-0 | 0.976570307 | 1 | 790 | 2296 | 52.5 | 0 | |
| In-0 | 0.676249637 | 2 | 960 | 1950 | 47.5 | 3 | |
| Is-0 | 0.66751979 | 1 | 830 | 2296 | 50.5 | 0 | |
| Ka-0 | 0.885133354 | 1.5 | 1332 | 2421 | 46.5 | 0 | |
| Kas-1 | 0.629291211 | 3 | 593 | 2354 | 35.0 | 0 | |
| Kavlinge-1 | 0.658837481 | 3 | 727 | 2318 | 55.8 | 0 | |
| Kelsterbach- | 0.521977785 | 1 | 756 | 2457 | 50.0 | 0 | |
| Kent | 0.828146853 | 2 | 728 | 2132 | 51.2 | 0 | |
| Kin-0 | 0.605019679 | 2 | 982 | 2472 | 44.5 | 3 | |
| Kn-0 | 0.946659376 | 1 | 725 | 2296 | 54.5 | 0 | |
| Kni-1 | 0.498661266 | 3 | 757 | 2266 | 55.7 | 0 | |
| Knox-10 | 0.633171897 | 3.5 | 1061 | 2786 | 41.3 | 0 | |
| Knox-18 | 0.432482616 | 3 | 1061 | 2786 | 41.3 | 0 | |

| | | | | | | |
|---|---|---|---|---|---|---|
| Koeln (PHW3 | 0.845433332 | 2 | 800 | 1595 | 50.9 | 3 |
| Kondara | 0.419948582 | 2 | 846 | 3477 | 38.5 | 0 |
| Kz-1 | 0.530563404 | 2 | 382 | 2528 | 49.5 | 0 |
| Kz-13 | 0.367846115 | 3 | 382 | 2528 | 49.5 | 0 |
| Kz-9 | 0.518829715 | 1 | 382 | 2528 | 49.5 | 0 |
| Lc-0 | 0.243585721 | 1.5 | 1073 | 1512 | 57.5 | 0 |
| Ler-1 | 0.56567497 | 2 | 1170 | 2625 | 48.0 | 1 |
| Li2:1 | 0.711160785 | 2 | 732 | 2251 | 50.4 | 0 |
| Liarum | 0.692624412 | 3 | 789 | 2214 | 55.9 | 0 |
| Lillo-1 | 0.516327471 | 3 | 664 | 2652 | 56.2 | 0 |
| Lip-0 | 0.628961649 | 1.5 | 836 | 2436 | 50.0 | 0 |
| Lis-1 | 0.677346545 | 3 | 767 | 2488 | 56.0 | 0 |
| Lis-2 | 0.819903212 | 3.5 | 767 | 2488 | 56.0 | 3 |
| Lisse (PHW-3 | 0.925749594 | 3.5 | 940 | 2476 | 52.3 | 3 |
| Ll-0 | 0.678830831 | 3 | 590 | 3242 | 41.6 | 3 |
| Lm-2 | 0.849735184 | 1 | 730 | 2275 | 48.0 | 3 |
| Lom1-1 | 0.948051491 | 3.5 | 786 | 2263 | 56.1 | 0 |
| Lov-1 | 0.551961345 | 3.5 | 685 | 2482 | 62.8 | 0 |
| Lov-5 | 0.785517267 | 3.5 | 685 | 2482 | 62.8 | 1 |
| Lp2-2 | 0.752288093 | 2 | 510 | 1840 | 49.4 | 3 |
| Lp2-6 | 0.78065811 | 2 | 510 | 1840 | 49.4 | 1 |
| Lu-1 | 0.899353765 | 3.5 | 757 | 2266 | 55.5 | 0 |
| Lund | 0.660133593 | 3.5 | 757 | 2266 | 55.7 | 0 |
| Lz-0 | 0.799192623 | 2 | 971 | 2454 | 46.0 | 0 |
| Mir-0 | 0.708220486 | 3 | 1559 | 3050 | 45.7 | 0 |
| Mr-0 | 0.93129799 | 3.5 | 1387 | 2804 | 44.2 | 0 |
| Mrk-0 | 0.871535787 | 2 | 910 | 2527 | 49.0 | 0 |
| Ms-0 | 0.696232405 | 3.5 | 678 | 2306 | 55.8 | 0 |
| Mt-0 | 0.674369376 | 2 | 98 | 3674 | 32.3 | 0 |
| Mz-0 | 0.684702424 | 2 | 1115 | 2649 | 50.3 | 3 |
| N13 | 0.69066023 | 3 | 766 | 2296 | 61.4 | 0 |
| Na-1 | 0.465385682 | 1.5 | 748 | 2373 | 47.5 | 0 |
| NC-6 | 0.618681868 | 3 | 1217 | 3023 | 35.1 | 0 |
| Nd-1 | 0.81691123 | 2 | 954 | 2306 | 50.3 | 0 |
| NFA-10 | 0.783529234 | 2 | 452 | 2199 | 51.4 | 3 |
| NFA-8 | 0.547607561 | 2 | 452 | 2199 | 51.4 | 3 |
| No-0 | 0.520303265 | 1 | 901 | 2458 | 51.0 | 0 |
| Nok-3 | 0.467427584 | 3.5 | 940 | 2476 | 52.2 | 0 |
| Nw-0 | 0.761458544 | 2 | 803 | 2275 | 50.5 | 0 |
| Omo2-3 | 0.951035051 | 3.5 | 664 | 2652 | 56.1 | 0 |
| Or-1 | 0.541408494 | 3.5 | 619 | 2388 | 56.5 | 0 |
| Ost-0 | 0.829072301 | 3.5 | 632 | 2503 | 60.5 | 0 |
| Ove-0 | 0.509906305 | 1.5 | 830 | 2266 | 53.3 | 3 |
| Oy-0 | 0.606994083 | 2 | 2646 | 1892 | 60.4 | 0 |
| Pa-1 | 0.430139904 | 1.5 | 647 | 3166 | 38.0 | 3 |
| Per-1 | 0.326010485 | 1.5 | 740 | 2208 | 58.0 | 0 |
| PHW-2 | 0.909314783 | 3 | 935 | 2923 | 43.7 | 0 |
| PHW-34 | 0.984360658 | 3.5 | 696 | 2351 | 48.6 | 0 |
| Pi-0 | 0.780869931 | 2 | 1672 | 2500 | 47.3 | 0 |
| Pla-0 | 0.994694391 | 2.5 | 717 | 3242 | 41.5 | 3 |
| Pna-10 | 0.613637119 | 2 | 1078 | 2731 | 42.1 | 0 |
| Pna-17 | 0.955360056 | 2 | 1078 | 2731 | 42.1 | 1 |
| Pro-0 | 0.65633195 | 2 | 1275 | 2214 | 43.3 | 3 |
| Pu2-23 | 0.666765485 | 2 | 672 | 2522 | 49.4 | 0 |
| Pu2-7 | 0.272904786 | 2 | 672 | 2522 | 49.4 | 0 |
| Pu2-8 | 0.316837307 | 2.5 | 672 | 2522 | 49.4 | 0 |
| Ra-0 | 0.511987898 | 2 | 858 | 2525 | 46.0 | 0 |
| Rak-2 | 0.546130388 | 1.5 | 612 | 2665 | 49.1 | 0 |
| Rd-0 | 0.667389053 | 1 | 803 | 2409 | 50.5 | 0 |
| Ren-1 | 0.743065968 | 2 | 737 | 2233 | 48.5 | 3 |
| Rev-1 | 0.79993819 | 3.5 | 770 | 2254 | 55.7 | 0 |
| Rmx-A02 | 0.660614753 | 2 | 1078 | 2731 | 42.0 | 3 |
| RRS-10 | 0.572710244 | 3.5 | 1078 | 2738 | 41.6 | 0 |
| RRS-7 | 0.746126358 | 3.5 | 1078 | 2738 | 41.6 | 0 |
| Rsch-4 | 0.551496645 | 1 | 661 | 2257 | 56.5 | 0 |
| Rubezhnoe-1 | 0.799038406 | 2 | 540 | 3559 | 49.1 | 0 |
| Sanna-2 | 0.630165734 | 3.5 | 736 | 2563 | 62.7 | 0 |
| Sap-0 | 0.699344217 | 1.5 | 672 | 1740 | 49.5 | 0 |
| Sav-0 | 0.460728837 | 1 | 510 | 1840 | 49.2 | 0 |
| Se-0 | 0.565953856 | 2 | 493 | 3099 | 38.3 | 0 |
| Sf-1 | 0.586579802 | 3 | 590 | 3242 | 41.8 | 3 |
| Sakhdara | 0.434108649 | 2 | 251 | 3328 | 39.0 | 0 |
| Sorbo | 0.421855766 | 3 | 280 | 3424 | 38.4 | 0 |

| | | | | | | |
|---|---|---|---|---|---|---|
| Spr1-2 | 0.95020318 | 3.5 | 325 | 2427 | 56.3 | 3 |
| Spr1-6 | 0.838332293 | 3.5 | 325 | 2427 | 56.3 | 3 |
| Sq-1 | 0.955170455 | 2 | 452 | 2199 | 51.4 | 3 |
| Sq-8 | 0.693134781 | 2 | 452 | 2199 | 51.4 | 0 |
| St-0 | 0.742235246 | 2 | 546 | 1898 | 59.0 | 0 |
| Stw-0 | 0.445468926 | 2.5 | 690 | 2366 | 52.5 | 0 |
| Ta-0 | 0.441769348 | 2 | 771 | 2412 | 49.5 | 0 |
| Tamm-2 | 0.438615733 | 3.5 | 678 | 2454 | 60.0 | 0 |
| Tamm-27 | 0.537765744 | 3.5 | 678 | 2454 | 60.0 | 0 |
| Tottarp-2 | 0.910679163 | 3.5 | 781 | 2157 | 56.3 | 3 |
| Ts-1 | 0.638200703 | 2 | 590 | 3242 | 41.7 | 0 |
| Ts-5 | 0.741321102 | 2 | 590 | 3242 | 41.7 | 3 |
| Tsu-0 | 0.682346629 | 1.5 | 2015 | 2832 | 34.4 | 0 |
| Tsu-1 | 0.619414839 | 1.5 | 2015 | 2832 | 34.4 | 0 |
| Tu-0 | 0.931319167 | 2.5 | 1002 | 3029 | 44.9 | 0 |
| Uk-3 | 0.976147508 | 3 | 1115 | 2649 | 48.1 | 0 |
| Ull2-3 | 0.580559475 | 2 | 786 | 2263 | 56.1 | 0 |
| Ull2-5 | 0.217612721 | 3 | 786 | 2263 | 56.1 | 0 |
| Uod-1 | 0.375045857 | 2 | 868 | 2603 | 48.3 | 0 |
| Uod-7 | 0.55087817 | 2 | 868 | 2603 | 48.3 | 0 |
| Van-0 | 0.649732575 | 2 | 2351 | 2530 | 49.5 | 0 |
| Var2-1 | 0.555156728 | 3.5 | 675 | 2518 | 55.6 | 0 |
| Var2-6 | 0.591419803 | 3.5 | 675 | 2518 | 55.6 | 0 |
| Vastervik | 0.368766451 | 3.5 | 647 | 2561 | 57.8 | 0 |
| Vimmerby | 0.605785731 | 3.5 | 696 | 2214 | 57.7 | 0 |
| Vinslov | 0.508388834 | 3.5 | 786 | 2263 | 56.1 | 0 |
| Wa-0 | 0.417995531 | 1 | 695 | 2461 | 52.3 | 0 |
| Wei-0 | 0.525062628 | 1 | 1544 | 2619 | 47.3 | 0 |
| Wil-1 | 0.474099904 | 1 | 764 | 2202 | 55.1 | 0 |
| Wil-2 | 0.376729822 | 1 | 764 | 2202 | 55.1 | 0 |
| Ws-0 | 0.558460597 | 2.5 | 610 | 2595 | 52.3 | 0 |
| Ws-2 | 0.68686151 | 1 | 610 | 2595 | 52.3 | 0 |
| Wt-5 | 0.698128756 | 2 | 812 | 2269 | 52.3 | 0 |
| Wu-0 | 0.945628195 | 2.5 | 757 | 2464 | 49.8 | 3 |
| Yo-0 | 0.569396169 | 2 | 567 | 3531 | 37.5 | 0 |
| Zdr-1 | 0.590762034 | 2 | 806 | 2324 | 49.4 | 0 |
| Zdr-6 | 0.390700247 | 2 | 806 | 2324 | 49.4 | 0 |
| Zu-1 | 0.638286729 | 1.5 | 1463 | 2699 | 47.4 | 0 |

### Supplementary Table 2

| purpose | Gene name | Gene number | sequence 5' - 3' | sequence 3' - 5' |
|---|---|---|---|---|
| QRT-PCR | ATG1a | At3g61960 | GAAACAAGTGCTGCCACTCA | GCTGCACTTTCTTTGTTACTCAAAT |
| QRT-PCR | ATG1b | At3g53930 | TGCCAAGTTTGTTGAATCTGA | CTCTGCGTCTCCCATCACTT |
| QRT-PCR | ATG1c | At2g37840 | TACTGCGCCGTAATCCAGT | CGAAAGAAAACCGTCCATTG |
| QRT-PCR | ATG8c | At1g62040 | TTTCAAGTTGGAACACCCACTA | AGCTCTCTCTACGATCACTGGAA |
| QRT-PCR | ATG8e | At2g45170 | ACCCTGATCGAATTCCTGTG | TTAGGTCTGATGGCACAAGGT |
| QRT-PCR | ATG8f | At4g16520 | TCCTGATAGGATTCCGGTGA | AAACTGCCCCACAGTCAGAT |
| QRT-PCR | ATG11 | At4g30790 | GCTTCTCGAAGAATCCCAGA | GATCAGCAGCACAAAGATGG |
| QRT-PCR | ATG13a | At3g49590 | GATGAGTCAGGACTTCAGTACAGC | TGTGAACCAGAACACCAACC |
| QRT-PCR | ATG13b | At3g18770 | CTTGCCCATTTGATGTTGAG | GGGTATGACCCGCTTGAAT |
| QRT-PCR | ATG101 | At5g66930 | TGGGAACAATGGTACATCAATCT | CTCTCCTCCGATGCCTCTC |
| QRT-PCR | GAPDH | At1g13440 | TTGGTGACAACAGGTCAAGCA | AAACTTGTCGCTCAATGCAATC |
| QRT-PCR | YLS8 | At5g08290 | TTGGTGATTGCTCCAAAAGA | AGTGTTGGGAAGCTCGATTAGT |
| Golden Gate | ATG101prom | At5g66930 | atatGGTCTCAGCGGGAGCTTGTTTGGTCCATGAG | atatGGTCTCTCAGATTCTTCGATATTTCTCTTTATCCTATTGG |
| Golden Gate | ATG101 gene | At5g66930 | atatGGTCTCATCTGAACAATGATGAAGCAAGCTTATGAAAAAG | atatGGTCTCTGGTGCCGCCGAGCATTGATGGATGG |
| SDM (SNP1) | ATG101 | At5g66930 | CTTGACATGCTcTCTGTATCCACTTTCGTTG | CAACGAAAGTGGATACAGAgAGCATGTCAAG |
| SDM (SNP3) | ATG101 | At5g66930 | GAAGTAATGATCTaATCCTTTTCATTGTCTCATCC | GGATGAGACAATGAAAAGGATtAGATCATTACTTC |

All genes are listed with their accepted name and accession numbers
